# Tissue-Resident and Effector-Memory Lymphocyte Recruitment to the Ocular Mucosa Requires CCL28/CCR10 Signaling During Recurrent HSV-1 Infection

**DOI:** 10.64898/2026.09.14.751393

**Authors:** Sweta Karan, Swayam Prakash, Chhaya Maurya, Sarah Xue LeNg, Lbachir BenMohamed

## Abstract

Ocular herpes simplex virus type 1 (HSV-1) infection is a major cause of infectious corneal disease that can lead to recurrent herpetic keratitis, chronic inflammation, and vision loss. Although effective antiviral immunity is essential for controlling HSV-1 and limiting recurrent disease, the mechanisms regulating recruitment of protective lymphocytes to the ocular mucosa remain incompletely understood. CCL28 is a mucosa-associated chemokine expressed at epithelial surfaces that signals through its receptor CCR10, expressed by distinct antiviral T-and B-cell subsets. Using a mouse model of recurrent ocular HSV-1 infection, we investigated the role of the CCL28/CCR10 axis in mobilizing protective HSV-1-specific immune cells to the ocular mucosa (OM). Following infection, wild-type (WT) B6 mice showed a marked increase in CCL28 expression within the OM, coinciding with accumulation of HSV-specific effector-memory (CCR10^+^CD44^+^CD62L^−^) T cells and CCR10^+^B220^+^ B cells in the OM and trigeminal ganglia (TG). These cells displayed an antigen-experienced, effector-memory phenotype, suggesting that CCL28/CCR10 signaling contributes to a localized antiviral immune response in the ocular mucosa. By contrast, CCL28-deficient (CCL28^(−/−)^) mice: (*i*) were more susceptible to recurrent ocular HSV-1 infection; (*ii*) exhibited significantly reduced frequencies of HSV-specific effector-memory (CCR10^+^CD44^+^CD62L^−^) and tissue-resident memory (CCR10^+^CD69^+^CD103^+^) T cells, as well as CCR10^+^B220^+^ B cells, in the infected OM and TG; and (*iii*) showed diminished effector function (IFN-γ, TNFα, and Granzyme B production) in these populations, associated with increased viral burden and ocular pathology. These findings identify the CCL28/CCR10 axis as a critical regulator of antiviral B-and T-cell trafficking to the ocular mucosa, contributing to protection against recurrent herpetic eye disease and providing a mechanistic basis for targeting mucosal chemokine pathways to improve vaccine strategies against ocular herpes.

**IMPORTANCE:** Herpes eye infections are a leading cause of blindness caused by an infectious agent. The virus that causes this disease, herpes simplex virus, can hide in nerve cells near the eye and reawaken periodically, causing repeated bouts of corneal damage. Our bodies rely on immune cells traveling to the eye’s surface to fight off these flare-ups, but how those cells know to travel there has not been well understood. This study shows that the CCL28 signaling molecule made by the surface tissue of the eye acts like a homing beacon, calling in specialized immune cells trained to fight this particular virus. Using mice, we found that animals lacking this CCL28 molecule struggled to control the initial infection and were far more likely to suffer damage when the virus reactivated. These animals also had fewer of the specialized defender cells present at the eye, and the cells they did have were weaker at fighting the virus. Our findings reveal a key piece of the body’s immune defense system against recurrent eye herpes and point to new ways this pathway could be harnessed to design better vaccines or treatments aimed at preventing the recurrent blinding damage caused by repeated viral flare-ups.

## INTRODUCTION

Herpes simplex virus type 1 (HSV-1) is a widespread human pathogen and a leading infectious cause of corneal blindness worldwide, with an estimated 3.72 billion new cases of HSV keratitis occurring globally each year (1–4). Ocular HSV-1 infection can cause epithelial and stromal keratitis, and recurrent episodes of infection and inflammation can result in progressive corneal scarring, neovascularization, vision impairment, and, in severe cases, blindness; approximately 230,000 people acquire new uniocular vision impairment from HSV keratitis each year (5, 6), and in the United States alone, roughly 500,000 individuals are affected by ocular HSV, with about one in five progressing to stromal keratitis and its attendant risk of blindness (7). Following primary infection, HSV-1 establishes lifelong latency in the sensory neurons of the trigeminal ganglia (TG) and can reactivate periodically, allowing the virus to traffic back to the cornea and cause recurrent disease, with each successive reactivation episode increasing the cumulative risk of vision-threatening stromal disease (8, 9). Although antiviral drugs can reduce viral replication and clinical disease, they do not eliminate latent virus or provide long-term protective immunity, and despite decades of research, an effective vaccine capable of preventing ocular HSV-1 infection or recurrent disease remains an unmet medical need (10, 11).

The outcome of ocular HSV-1 infection is determined by a complex interaction between viral replication and the host immune response (12). The corneal epithelial cells and infiltrating immune cells produce a variety of cytokines and chemokines to regulate the recruitment of innate and adaptive immune cells to the infected tissue (13, 14). Neutrophils, monocytes, NK cells, dendritic cells, B cells, and T cells all contribute to the local response to HSV-1 (15, 16). While these inflammatory responses are necessary for controlling viral replication, excessive or poorly regulated inflammation can also contribute to corneal pathology (17). Thus, effective immunity against ocular HSV-1 requires not only the generation of antiviral lymphocytes but also their appropriate localization and maintenance at the site of infection (18).

T cells play a central role in controlling HSV-1 infection. HSV-specific T cells migrate to peripheral tissues where they can recognize and eliminate infected cells and provide local immune surveillance (16). Memory CD8^+^ T cells are particularly important for controlling HSV-1 at the site of infection and during viral reactivation (19). B cells and antibody responses also contribute to antiviral protection by neutralizing extracellular virus and limiting viral dissemination (20). However, the mechanisms that determine how HSV-specific memory B and T cells are recruited and maintained within the ocular tissue remain incompletely understood.

Chemokines are important regulators of trafficking of immune cells between the circulation and peripheral tissues (21). Among the roughly 48 chemokines identified in humans, a small subset, CCL25, CCL28, CXCL14, and CXCL17, is distinguished by constitutive, homeostatic expression at mucosal epithelial surfaces rather than induction solely by inflammation (22). Several chemokines are induced during HSV-1 infection, but most studies have focused on inflammatory chemokines involved in the early recruitment of innate immune cells (23). The chemokine pathways responsible for regulating the localization of antigen-specific memory lymphocytes at the ocular site of HSV-1 infection has not been explored. Identifying these pathways is particularly important for vaccine development because durable protection may depend on establishing a pool of antiviral memory cells capable of responding rapidly at the site where HSV-1 replicates and reactivates.

CCL28 is a mucosa-associated chemokine that is expressed by epithelial tissues and signals primarily through the chemokine receptor CCR10 (24). The CCL28/CCR10 axis has been implicated in the trafficking of specific B-and T-cell populations to mucosal tissues (25, 26). Due to the importance of local memory lymphocytes in antiviral protection, we hypothesized that CCL28 may contribute to the recruitment of protective HSV-1-specific immune cells to ocular tissues. However, the role of the CCL28/CCR10 pathway in ocular HSV-1 infection and vaccine-induced immunity remains unclear.

In this study, we investigated the role of the CCL28/CCR10 axis in regulating local antiviral immunity following ocular HSV-1 infection. We examined CCL28 expression and the accumulation of CCR10-expressing immune cells in ocular tissues and characterized HSV-1-specific memory T-and B-cell responses following infection. We further evaluated CCL28 contribution to antiviral immunity using CCL28-deficient (CCL28^(−/−)^) mice. Our findings show that ocular HSV-1 infection is associated with increased CCL28 expression and accumulation of CCR10-expressing antiviral lymphocytes at the site of infection. Loss of CCL28 resulted in impaired recruitment of these immune populations and compromised local antiviral immunity. Together, our results suggest that the CCL28/CCR10 axis is an important component of the local immune response to ocular HSV-1 and provide a rationale for exploiting this pathway to improve vaccine-induced mucosal immunity against ocular herpes.

## RESULTS

### 1. HSV-1 induces CCL28 expression in primary human corneal epithelial cells

Infection of primary human corneal epithelial cells (primary HCECs) with HSV-1 at the multiplicity of infection (MOI) of 2.5 led to progressive cytopathic changes starting around 6 – 48 h post-infection, characterized by prominent cellular rounding, loss of adherence, and cell lysis by 24 – 48 h compared to the confluent monolayer of uninfected cells (**Fig. 1A**). The transcriptional response and expression levels of mucosal and homeostatic chemokines to viral infection in primary human corneal epithelial cells (HCECs) were evaluated over a 48-hour time course. Under uninfected conditions, HCECs exhibited differential relative transcript levels among the target chemokines (ΔCt relative to housekeeping controls), with CXCL14 showing the highest baseline abundance (∼−2.2), followed by moderate levels of CXCL17 (∼−4) and CCL28 (∼−4), and minimal expression of CCL25 (∼−7.1) (**Fig. 1B**). Following HSV-1 infection, the chemokines displayed distinct kinetic profiles. Expression levels of CXCL17, CXCL14, and CCL28 remained near baseline during the early phase of infection (2–6 h post-infection) while the expression of *CCL25* was not observed (**Fig. 1C**). However, CCL28 and CXCL17 chemokines showed significant upregulation from 12 h post-infection, accelerating continuously above baseline by 48 h (**Fig. 1C**). In contrast, CXCL14 expression was rapidly suppressed beyond 12 h post-infection and remained decreased through 48 h. These data indicate that HSV-1 infection in primary corneal epithelial cells induces a robust transcriptional activation of CCL28 and CXCL17, while concurrently repressing CXCL14 baseline expression. To determine whether the transcriptional upregulation of CCL28 translated to increased protein expression, primary human corneal epithelial cells (HCECs) were evaluated via immunofluorescence microscopy following HSV-1 infection. HSV-1-infected primary HCECs demonstrated significant immunoreactivity for CCL28 protein (red fluorescence, white arrows) (**Fig. 1D**). CCL28 protein accumulated predominantly within the cytoplasm and perinuclear regions of infected corneal epithelial cells, displaying prominent localization surrounding the DAPI-stained nuclei (blue) (**Fig. 1D**). These results validate our qPCR findings, confirming that HSV-1 infection stimulates strong translational expression and cellular accumulation of the homeostatic chemokine CCL28 in primary human corneal epithelial cells.

**Figure 1.**
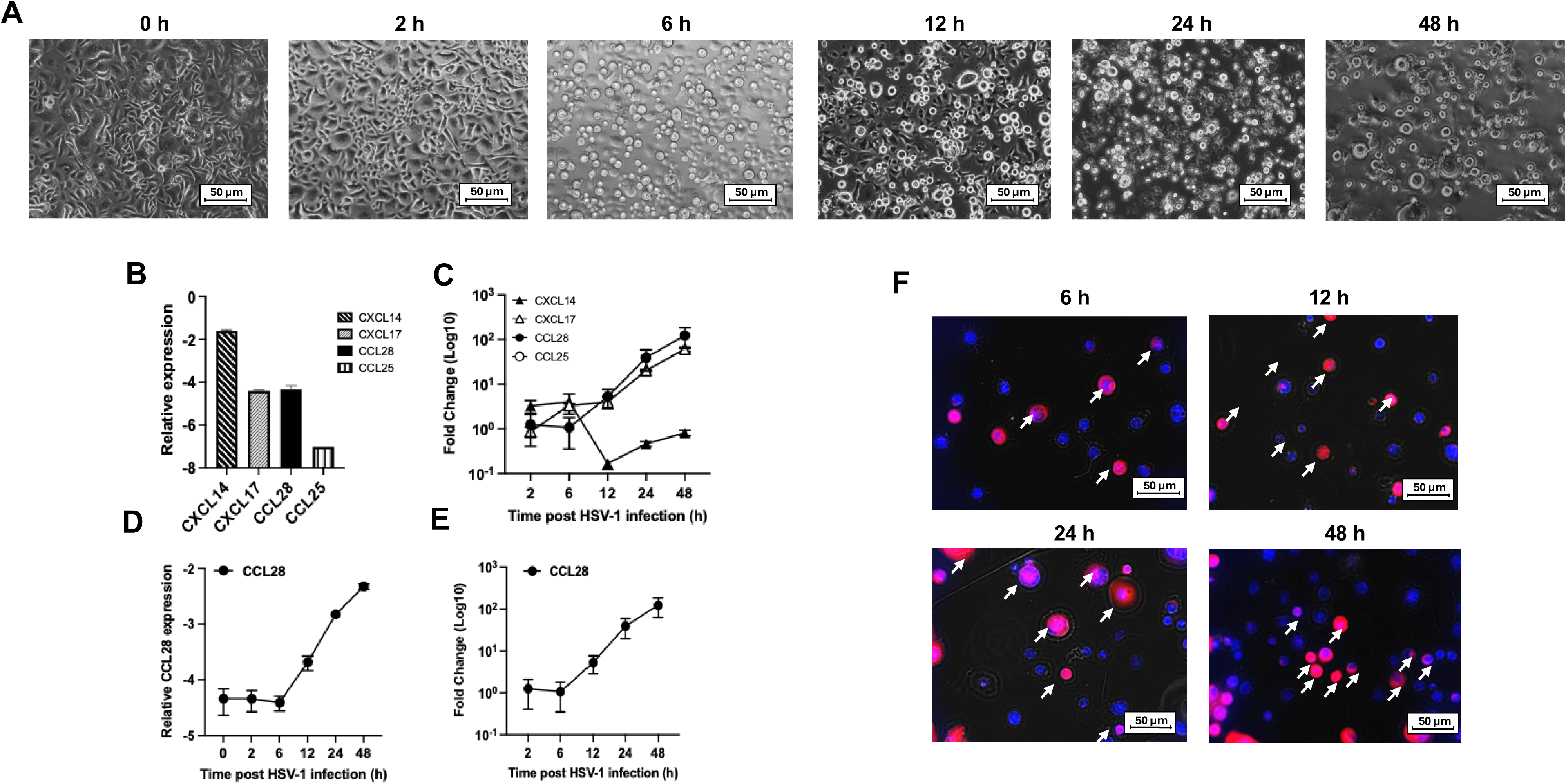
HSV-1 induces CCL28 expression in primary human corneal epithelial cells (HCECs): (**A**) Representative phase-contrast microscopy images showing cytopathic effects and morphological changes in HSV-1-infected HCECs over time (0 h, 6 h, 12 h, 24 h, and 48 h post-infection). Progressive cell rounding, detachment, and plaque formation are evident at later time points. Non-infected HCECs display a healthy monolayer epithelial morphology. (**B**) Baseline relative mRNA expression levels of the mucosal chemokines CXCL14, CXCL17, CCL28, and CCL25 in uninfected primary human corneal epithelial cells (HCECs), expressed relative to the housekeeping gene GAPDH, as measured by RT-qPCR. (**C**) Time-course kinetics showing log10 fold changes in CXCL14, CXCL17, CCL28, and CCL25 transcript levels across a 48-hour infection period (2 h, 6 h, 12 h, 24 h, and 48 h post-HSV-1 infection). (**D**) Absolute relative transcript levels (ΔCt) of CCL28 from 0 h to 48 h post-infection, demonstrating stable baseline levels in early infection followed by a persistent increase from 12 h onwards. (**E**) Quantification of CCL28 mRNA fold change (log10 scale) relative to baseline control (0 h or uninfected control), displaying multi-log induction at 48 h post-infection. Data are represented as mean ± SD from representative independent experiments. (**F**) Representative immunofluorescence images showing CCL28 protein localization and expression in primary human corneal epithelial cells (HCECs) following HSV-1 infection. Red fluorescence indicates CCL28 protein accumulation (white arrows point to CCL28-positive cells). Nuclei are counterstained with DAPI (*blue*).

### 2. UV-B light induces latent HSV-1 reactivation

To model recurrent herpes simplex virus type 1 (HSV-1) keratitis, latently infected mice were subjected to UV-B irradiation at 21 days post-primary infection. UV-B exposure successfully induced viral reactivation and subsequent recurrent herpes stromal keratitis (HSK) in both CCL28^(⁻/⁻)^ deficient and WT mice. Fluorescein staining of the cornea indicated corneal ulceration and severe herpetic keratitis in CCL28^(⁻/⁻)^ deficient compared with WT mice (**Fig. 2B**). Clinical scoring revealed that corneal pathology peaked between days 5 and 10 post-reactivation. Compared with WT mice, CCL28^(⁻/⁻)^ deficient showed significantly more severe corneal opacity, neovascularization, and clouding (*P* = 0.0352) **(Fig. 2C)**. Viral genome shedding was quantified as HSV-1 DNA copy number from corneal swabs, which showed increased HSV-1 DNA copy number following UV-B exposure in both groups, with CCL28^(⁻/⁻)^ deficient showing significantly increased viral shedding on days 4 (*P* < 0.0001), 6 (*P* = 0.036), and 10 (*P* = 0.0182) compared with WT mice (**Fig. 2E**); this elevated shedding correlated with the more severe corneal lesions observed in CCL28^(⁻/⁻)^ deficient mice. Survival analysis also revealed a significant difference between the two groups, with higher mortality in CCL28^(⁻/⁻)^ deficient mice (*P* = 0.0342) (**Fig. 2D**).

**Figure 2.**
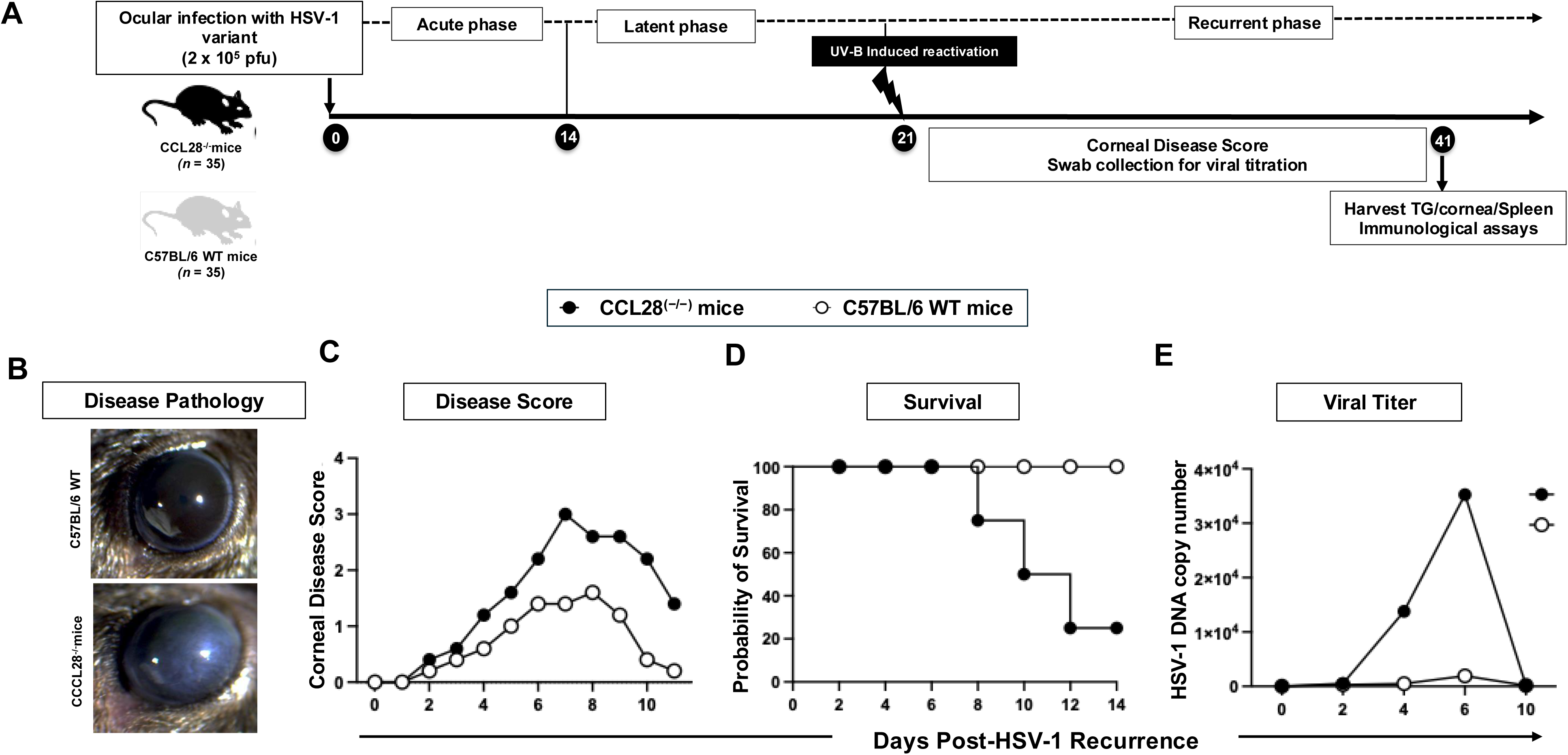
UV-B-induced HSV-1 recurrence, disease progression, and survival in CCL28^(−/−)^ deficient and Wild-Type (WT B6) mice. (**A**) Experimental timeline: CCL28^(−/−)^ deficient mice (*n* = 35) and WT B6 mice (*n* = 35) were ocularly infected with HSV-1 on day 0 to establish acute infection. Establishments of latency occurred by day 14 (14–21 days post-infection; dpi). At day 21, latent infection was reactivated using UV-B irradiation. Following reactivation, mice were monitored during the recurrent phase (21–41 dpi) for corneal disease scores and viral shedding via corneal swabs. Tissues (trigeminal ganglia (TG), cornea, and spleen) were harvested at day 41 for immunological assays. (**B**) Representative slit-lamp microscopy images display corneal disease pathology, showing mild stromal opacity (top) in WT B6 mice compared with severe corneal clouding and scarring (bottom) in CCL28^(−/−)^ deficient mice. (**C**) Corneal disease score: Disease scores peak during the recurrent phase from days 5–10 post-reactivation. (**D**) Kaplan-Meier survival analysis indicating the probability of survival following UV-B-induced viral reactivation across experimental groups. (**E**) Quantification of HSV-1 DNA copy numbers measured from corneal swab samples by qPCR across days post-reactivation demonstrates the time course of viral shedding.

### 3. CCL28 deficiency significantly impairs immune cell infiltration

To assess the role of CCL28 in immune cell homing during ocular infection, total immune cell populations were analyzed by multi-parameter flow cytometry in the trigeminal ganglia (TG) and corneas of CCL28^(-/-)^ deficient and WT mice. CCL28^(-/-)^ deficient mice displayed significantly reduced leukocyte populations compared to WT controls. We observed significantly lower frequencies of total CD45^+^ immune cell (*P* < 0.05) in CCL28^(−/−)^ deficient mice **(Fig. 3)**. This deficiency was consistent among B220^+^ B cells (*P* < 0.05), total CD3^+^ T cells (*P* < 0.05), CD4^+^ T cells (*P* < 0.01), and CD8^+^ T cells (*P* < 0.01) **(Fig. 3)**. We observed similar phenotype within the corneal tissue, where CCL28^(-/-)^ deficient mice exhibited significantly lower infiltration relative to WT mice. The infiltrating frequency of CD45^+^ total leukocytes was significantly decreased (*P* < 0.05) **(Fig. 3)**. Similar lower frequency of B220^+^ B cells (*P* < 0.01), CD3^+^ T cells (*P* < 0.05), CD4^+^ T cells (*P* < 0.05), and CD8^+^ T cells (*P* < 0.05) **(Fig. 3)** was also observed. These results demonstrate that CCL28 plays a critical role in mediating the homing and recruitment of adaptive lymphocyte populations (CD4^+^ T cells, CD8^+^ T cells, and B cells) and overall immune cell infiltration into both peripheral (cornea) and neural (trigeminal ganglia) sites during recurrent ocular infection.

**Figure 3.**
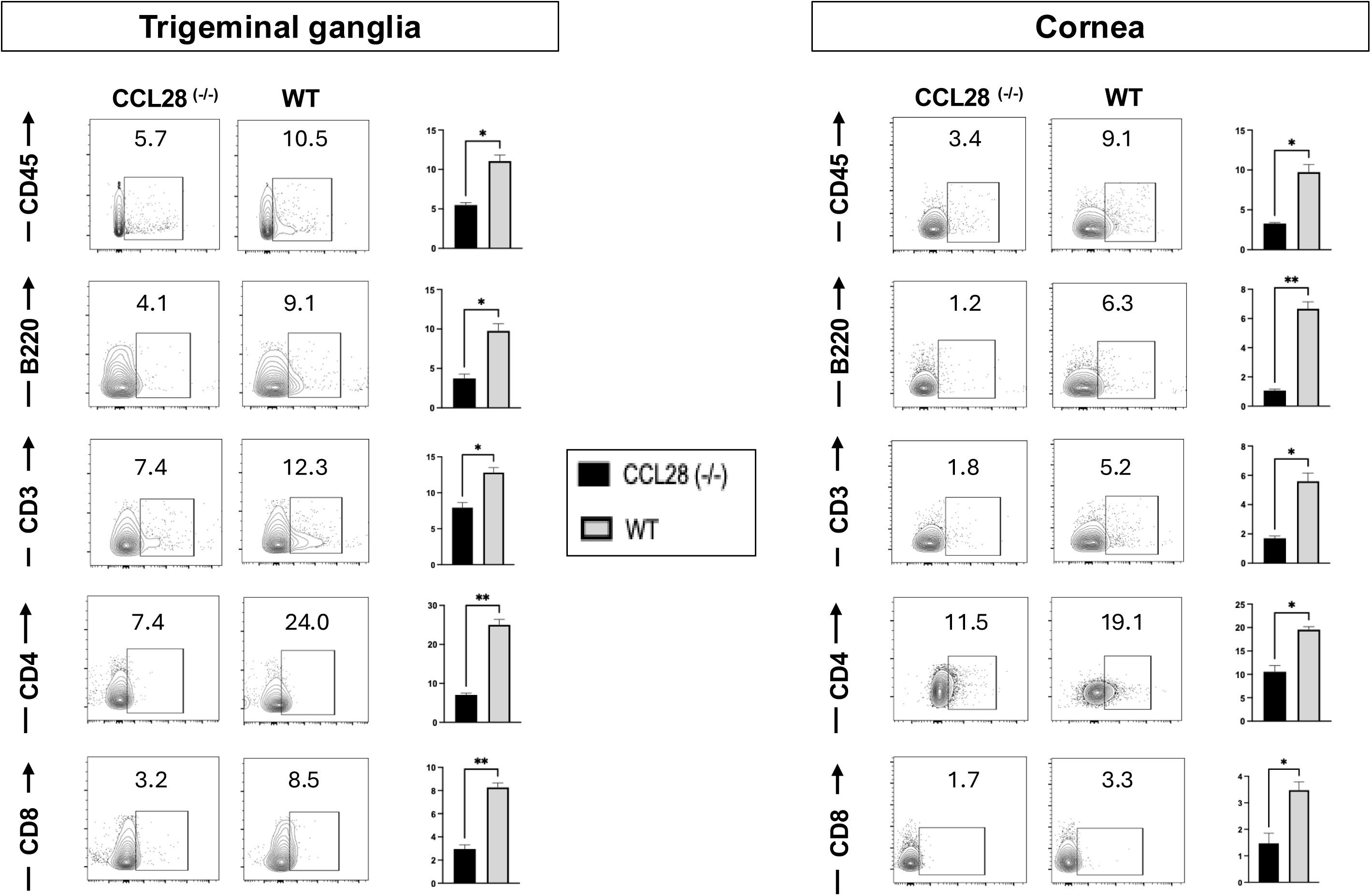
CCL28 deficiency impairs infiltration of immune cells in trigeminal ganglia and corneas of CCL28^(−/−)^ deficient mice. Representative contour plots display frequencies of total immune cells (CD45^+^), B cells (B220^+^), T cells (CD3^+^), and T cell subpopulations (CD4^+^ and CD8^+^) from single-cell suspensions of the trigeminal ganglia (TG, left panel) and corneas (right panel) of CCL28^(−/−)^ deficient (black bars) and WT (gray bars) mice. The graphs represent the frequency of each leukocyte subset per tissue. Data are represented as mean ± SD. Statistical significance between CCL28^(−/−)^ deficient and WT groups was assessed by an unpaired t-test (\**P* < 0.05, ** *P* < 0.01).

### 4. CCL28 deficiency reduces the frequency of infiltrating CCR10+ lymphocyte in cornea and TG

To determine whether loss of CCL28 impacts the recruitment of lymphocytes expressing its cognate receptor, flow cytometry was performed to quantify CCR10^+^ expression on B cells and T cell populations within the trigeminal ganglia (TG) and corneas of CCL28^(-/-)^ deficient and WT mice. The proportion of lymphocytes expressing CCR10^+^ was significantly reduced across all evaluated subsets in CCL28^(-/-)^ deficient mice compared to WT controls. Specifically, CCR10^+^ frequencies were decreased among B220**^+^** B cells (*P* < 0.01), total CD3**^+^** T cells (*P* < 0.01), CD4**^+^** T cells (*P* < 0.01), and CD8**^+^** T cells (*P* < 0.01) in TG **(Fig. 4)**. We observed similar reduction in CCR10^+^ cell frequencies within the corneal tissue of CCL28^(−/−)^ deficient mice relative to WT controls. Frequencies of CCR10^+^ expression decreased significantly across B220^+^ B cells (*P* < 0.01), CD3**^+^** T cells (*P* < 0.01), CD4**^+^** T cells (*P* < 0.01), and CD8**^+^** T cells (*P* < 0.05) **(Fig. 4)**. These data demonstrate that the absence of CCL28 severely limits the local accumulation and infiltration of CCR10^+^ B cells, CD4^+^ T cells, and CD8^+^ T cells at both peripheral ocular infection sites (cornea) and sensory neural tissues (trigeminal ganglia).

**Figure 4.**
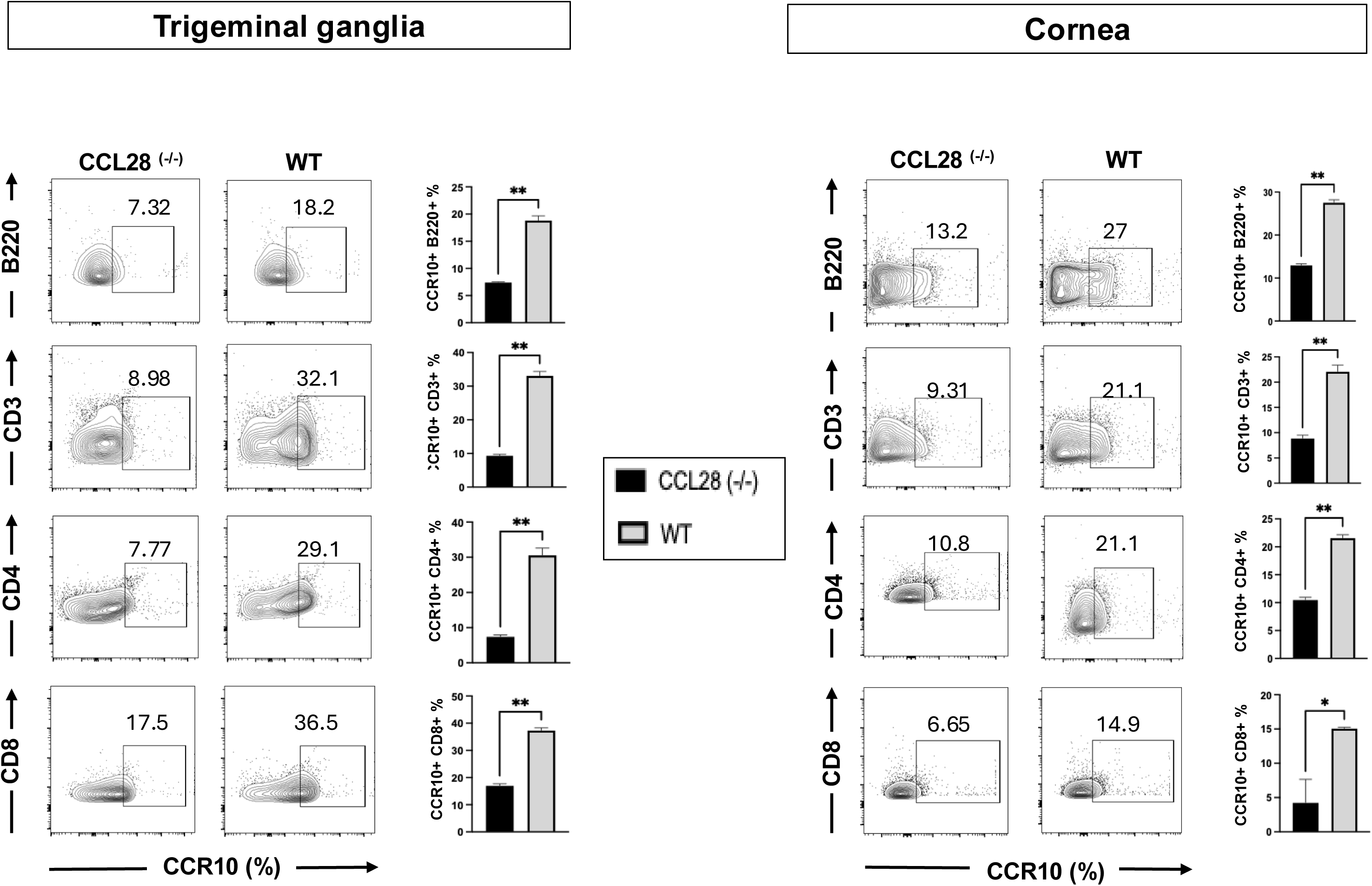
CCL28 deficiency reduces the frequency of infiltrating CCR10^+^ lymphocyte subsets in cornea and TG of CCL28^(−/−)^ deficient mice: Representative contour plots showing frequencies of CCR10-expressing B cells (B220^+^), total T cells (CD3^+^), and T cell subsets (CD4^+^ and CD8^+^) gated from single-cell suspensions isolated from the trigeminal ganglia (TG, left panel) and corneas (right panel) of CCL28^(−/−)^ deficient mice (black bars) and WT (gray bars) mice. The graphs represent the frequency of CCR10^+^ cells within each indicated lymphocyte lineage per tissue. Data represent mean ± SD. Statistical significance between CCL28^(−/−)^ deficient and WT groups was assessed by an unpaired t-test (* *P* < 0.05, ** *P* < 0.01).

### 5. CCL28 deficiency impairs effector memory CD4^+^ and CD8^+^ T cells

To evaluate the impact of CCL28 deficiency on T cell activation at sites of infection, expression of memory differentiation markers (CD44 and CD62L) was analyzed on infiltrating CD4^+^ and CD8^+^ T cell populations. Frequencies of effector memory (CD44^+^CD62L^−^) T cells were significantly lower in CCL28^(-/-)^ compared to WT controls. We observed reduced effector memory T cells (CD4^+^CD44^+^CD62L^−^) in CCL28^(−/−)^ TG compared to WT TG (*P* < 0.01). Similarly, effector memory (CD8^+^CD44^+^CD62L^−^) T cells decreased significantly in CCL28^(−/−)^ TG relative to WT TG (*P* < 0.01) (**Fig. 5**). A similar decline in effector memory T cell recruitment and retention occurred within the corneal of CCL28^(-/-)^ deficient mice. The frequency of CD4^+^CD44^+^CD62L^−^ T_EM_ cells was reduced in CCL28^(-/-)^ corneas compared to WT cornea *P* < 0.01). Likewise, CD8^+^CD44^+^CD62L^−^ T_EM_ cells were significantly reduced in CCL28^(-/-)^ cornea relative to WT cornea (*P* < 0.01) **(Fig. 5)**. The elevated proportion of T_EM_ cells in WT mice indicates effective recruitment and tissue retention of functional effector memory T cells at the HSV-1 infection site.

**Figure 5.**
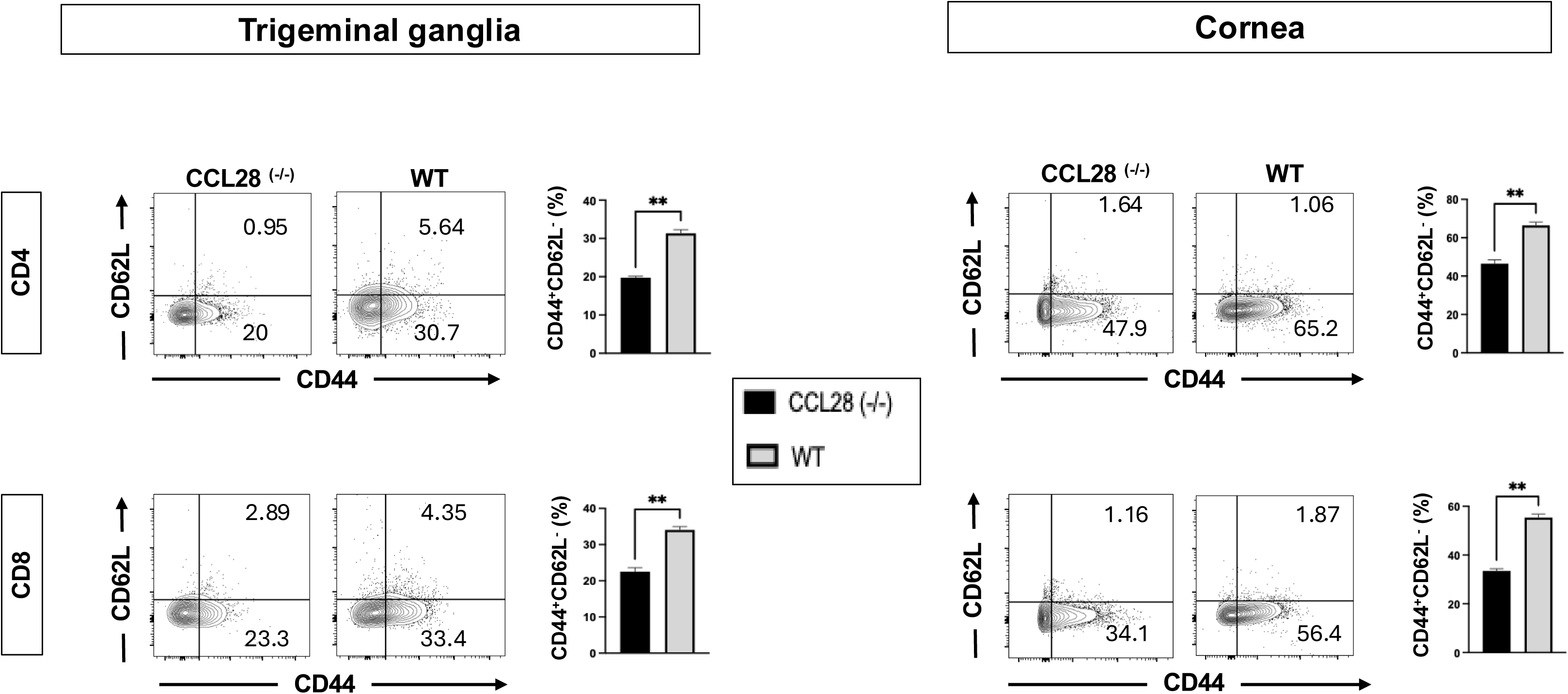
CCL28 deficiency impairs infiltration and retention of effector memory T cells (T_EM_): Representative flow cytometry contour plots showing gating for effector memory T cells (CD44^+^CD62L^−^) within CD4^+^ (top panel) and CD8^+^ (bottom panel) T cell populations isolated from the trigeminal ganglia (TG, left panel) and corneas (right panel) of CCL28^(-/-)^ deficient (black bars) and WT (gray bars) mice. The graphs represent the frequencies of effector memory CD4^+^ and CD8^+^ T cells (CD44^+^CD62L^−^) per tissue. Data represent mean ± SD. Statistical significance between CCL28^(−/−)^ deficient and WT groups was assessed by an unpaired t-test (** *P* < 0.01).

### 6. CCL28 deficiency impairs retention of tissue-resident memory (T_RM_) CD4^+^ and CD8^+^ T cells

To determine whether CCL28 regulates the establishment and maintenance of tissue-resident memory T cells at infection sites, expression of residence markers (CD69 and CD103) was evaluated on infiltrating CD4^+^ and CD8^+^ T cells. Frequencies of T cells expressing activation and retention markers were significantly diminished in CCL28^(-/-)^ compared to WT controls. We observed reduced expression of CD69^+^ in CCL28^(-/-)^ deficient mice derived TG across both CD4^+^ T cells (*P* < 0.01) and CD8^+^ T cells (*P* < 0.01) **(Fig. 6)**. Similarly, CD103^+^ expression was significantly reduced in CCL28^(-/-)^ deficient mice derived TG among CD4^+^ T cells (*P* < 0.01) and CD8^+^ T cells (*P* < 0.05) **(Fig. 6)**. We observed similar reduction in tissue-resident memory subsets in the cornea of CCL28^(-/-)^ deficient mice. Frequencies of CD69^+^ expressing cells were sharply reduced in CCL28^(-/-)^ deficient mice derived corneas relative to WT controls for both CD4^+^ T cells (** P < 0.01) and CD8^+^ T cells (*P* < 0.001) **(Fig. 6)**. Furthermore, CD103^+^ expression was reduced on CCL28^(-/-)^ deficient mice derived corneal CD4^+^ T cells (*P* < 0.05) and CD8^+^ T cells (*P* < 0.05) **(Fig. 6)**. The significantly increased expression of CD69 and CD103 on WT T cells demonstrates effective retention of functional T_RM_ cells at the HSV-1 infection site.

**Figure 6.**
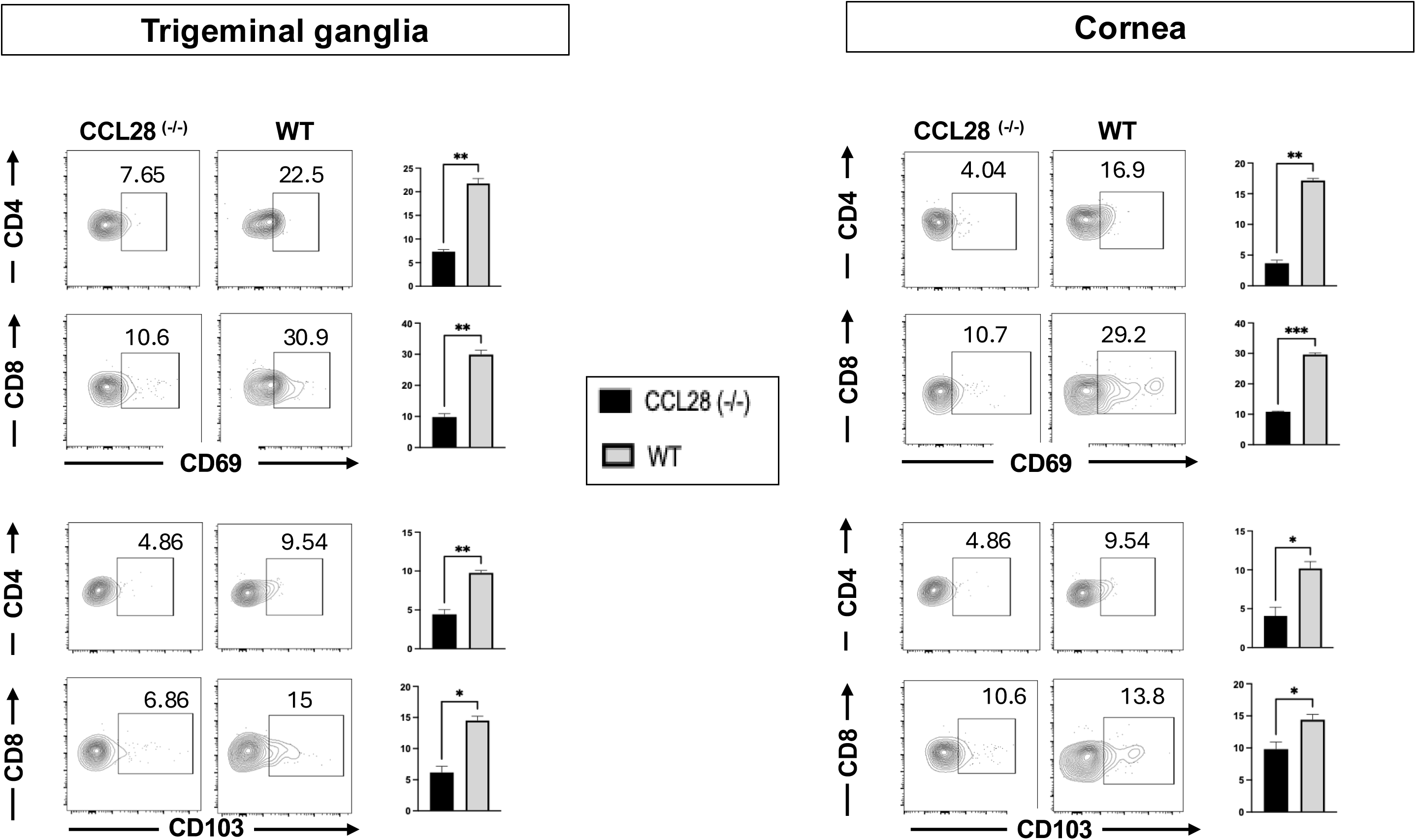
CCL28 deficiency impairs expression of tissue-resident memory T cell (T_RM_) Representative flow cytometry contour plots showing expression of tissue residence markers CD69 (top panels) and CD103 (bottom panels) on gated CD4^+^ and CD8^+^ T cell populations isolated from the trigeminal ganglia (TG, left) and corneas (right) of CCL28^(−/−)^ deficient (*black bars*) and WT (*gray bars*) mice. The graphs represent the frequencies of CD69^+^ or CD103^+^ subsets among CD4^+^ and CD8^+^ T cells per tissue. Data represent mean ± SD. Statistical significance between CCL28^(−/−)^ deficient and WT groups was assessed by an unpaired t-test (\**P* < 0.05, \*\**P* < 0.01, *** *P* < 0.001).

### 7. CCL28 deficiency impairs the functional effector response of CD4^+^ and CD8^+^ T cells

To evaluate whether CCL28 deficiency impacts the functional efficacy of infiltrating lymphocytes, intracellular production of cytotoxic (GzmB) and antiviral/inflammatory cytokines (IFN-γ and TNFα) was quantified in CD4^+^ and CD8^+^ T cells isolated from the trigeminal ganglia (TG) and cornea of CCL28^(-/-)^ deficient and WT mice. Frequencies of effector T cells were significantly reduced in CCL28^(-/-)^ deficient mice compared to WT controls. Intracellular GzmB^+^ levels were lower in CCL28^(-/-)^ TG for both CD4^+^ (*P* < 0.05) and CD8^+^ T cells (*P* < 0.05) **(Fig. 7)**. Similarly, IFN-γ^+^ frequencies were reduced in CD4^+^ (*P* < 0.05) and CD8^+^ T cells (*P* < 0.05) **(Fig. 7)**. We also observed decreased frequency of TNFα^+^ in CCL28^(- /-)^ deficient mice in CD4^+^ (*P* < 0.01) and CD8^+^ T cells (*P* < 0.01) **(Fig. 7)**. We also observed reduced T cell effector function in corneal tissue. Expression of GzmB^+^ was substantially decreased in CCL28^(-/-)^ deficient cornea relative to WT controls across CD4^+^ (*P* < 0.01) and CD8^+^ T cells (* P < 0.05) **(Fig. 7)**. Similarly, IFN-γ^+^ positive cells decreased in CD4^+^(*P* < 0.05) and CD8^+^ T cells (*P* < 0.05) **(Fig. 7)**. Finally, TNFα^+^ production was severely reduced in CCL28^(-/-)^ deficient mice corneal CD4^+^ (*P* < 0.05) and CD8^+^ T cells (*P* < 0.01) **(Fig. 7)**. These results showed that CCL28 deficiency severely reduces the proportion of functional, cytokine-producing, and cytotoxic CD4^+^ and CD8^+^ T cells in both peripheral (cornea) and sensory neural (TG) sites during HSV-1 infection.

**Figure 7.**
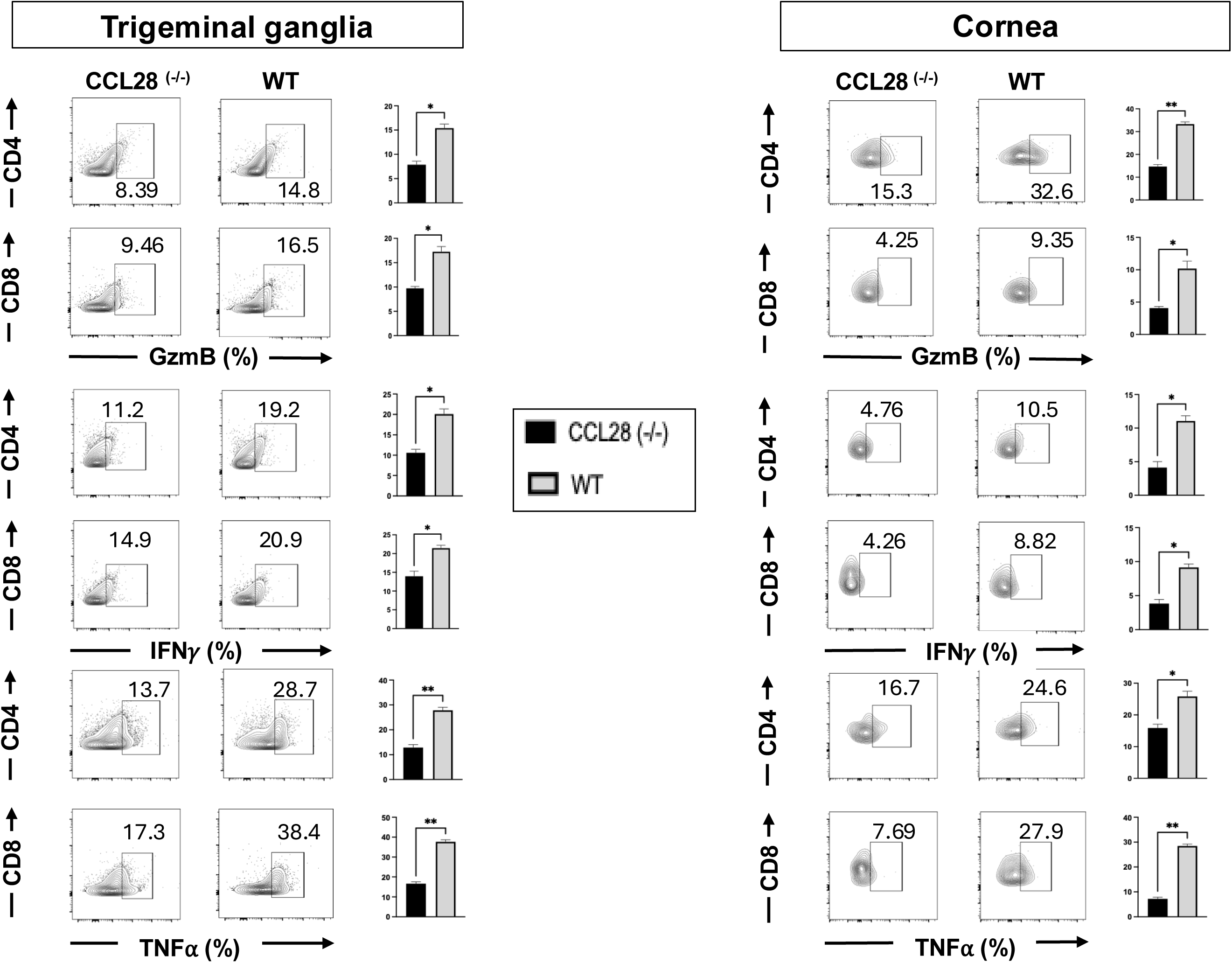
CCL28 deficiency impairs expression of cytotoxic and inflammatory effector function: Representative flow cytometry contour plots display intracellular expression of Granzyme B (GzmB), Interferon-gamma (IFN-γ), and Tumor Necrosis Factor-alpha (TNF-α) on gated CD4^+^ and CD8^+^ T cell populations isolated from the trigeminal ganglia (TG, left panel) and corneas (right panel) of CCL28^(−/−)^ deficient (black bars) and WT (gray bars) mice following HSV-1 infection. The graphs present the frequencies of GzmB^+^, IFN-γ^+^, and TNF-α^+^ expressing cells within CD4^+^ and CD8^+^ T cell compartments per tissue. Data represent mean ± SD. Statistical significance between CCL28^(−/−)^ deficient and WT groups was assessed by an unpaired t-test (* *P* < 0.05, ** *P* < 0.01).

### 8. CCL28 deficiency impairs the expression of CCR10 and infiltration of CD4^+^ and CD8^+^ T cells

To confirm in situ protein localization and immune cell infiltration within neural tissue, immunofluorescence staining of trigeminal ganglia (TG) tissue sections was evaluated in CCL28^(-/-)^ deficient and WT mice. WT trigeminal ganglia exhibited prominent, widespread expression of CCL28 protein (green, white arrows) and its cognate receptor CCR10 following infection **(Fig. 8)**. In contrast, CCL28^(-/-)^ deficient mice demonstrated a complete absence of CCL28 signal and a corresponding marked reduction in CCR10-expressing cells within the ganglia (**Fig. 8**). We observed reduced in situ infiltration of effector T lymphocytes in the TG of CCL28^(-/-)^ deficient mice compared to WT controls. CCL28^(-/-)^ deficient mice displayed reduced T cell signal, whereas WT mice showed dense clusters of CD4**^+^** and CD8**^+^** T cells (green, white arrows) **(Fig. 8)**. These observations confirm that CCL28 expression within the trigeminal ganglia is essential for recruiting CCR10**^+^** T cell subpopulations (CD4**^+^** and CD8**^+^**) into neural tissue following viral infection **(Fig. 8)**. Similarly, histological examination using H&E staining further demonstrated more extensive epithelial damage and inflammation in the absence of CCL28 (black arrows) **(Fig. 9)**. These histopathological observations demonstrate that CCL28 is essential for orchestrating robust leukocyte infiltration into both neural (trigeminal ganglia) and ocular tissues during HSV-1 infection.

**Figure 8.**
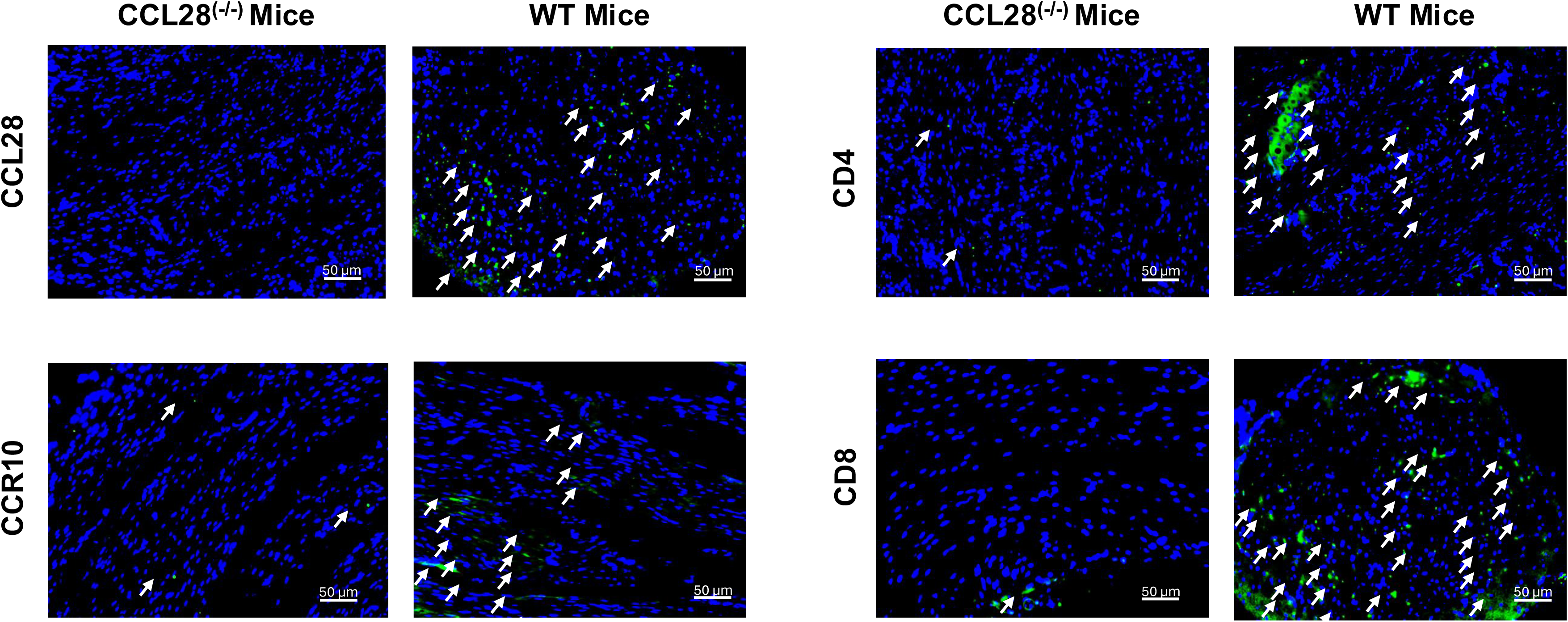
CCL28 deficiency impairs expression of CCL28, receptor CCR10, and Infiltrating CD4^+^ and CD8^+^ T cells: Representative immunofluorescence image displays expression of CCL28 (*top left*), its cognate receptor CCR10 (bottom left), infiltration of CD4 (top right), and CD8 (bottom right) in trigeminal ganglia (TG) tissue sections harvested from CCL28^(−/−)^ deficient and WT mice following HSV-1 infection. Green fluorescence indicates positive immunoreactivity for target proteins (CCL28, CCR10, CD4, or CD8, highlighted by white arrows). Cell nuclei are counterstained with DAPI (*blue*).

**Figure 9.**
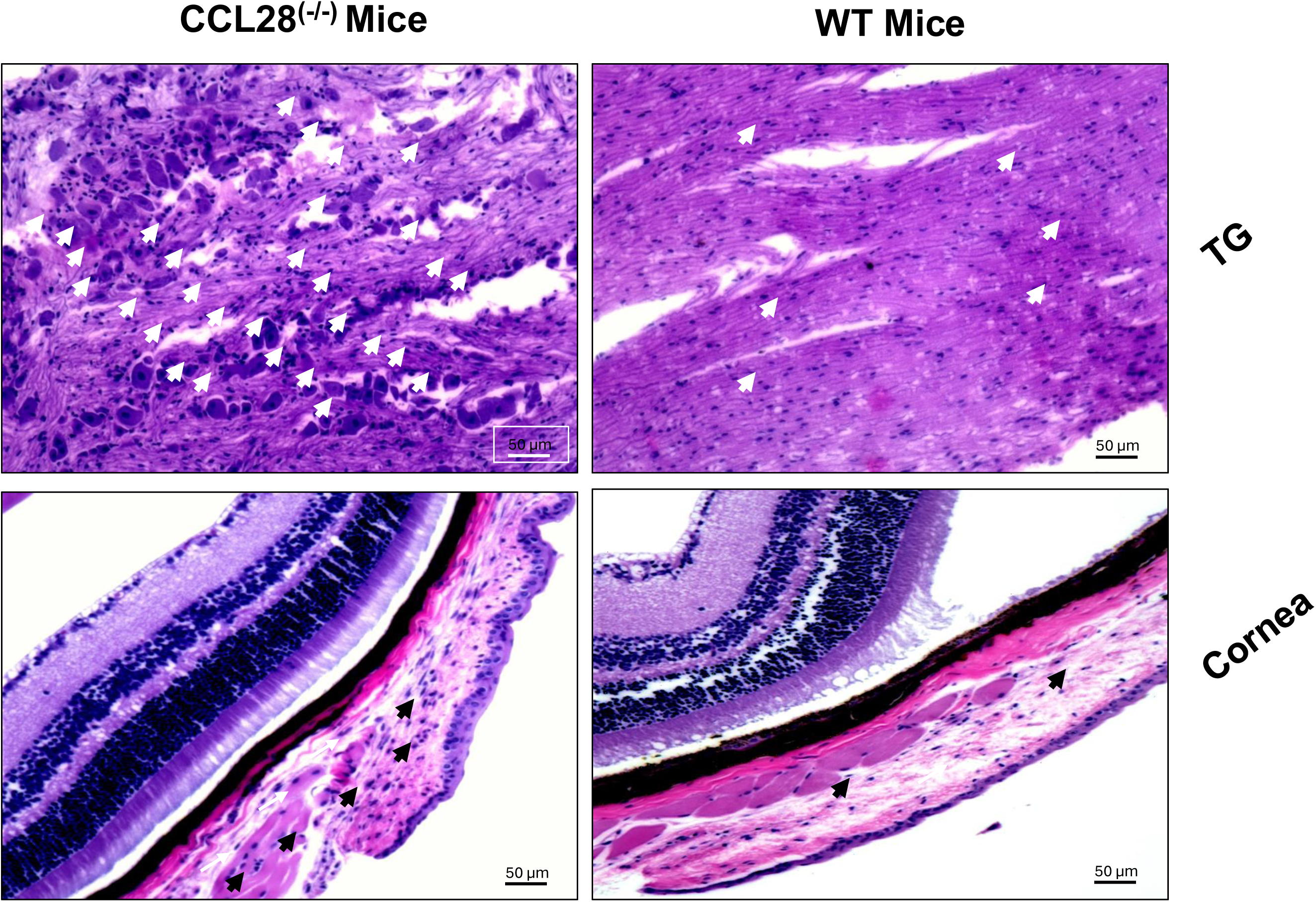
Histopathological evaluation of ocular pathology in CCL28^(−/−)^ deficient and WT mice: Representative Hematoxylin and Eosin (H&E)-stained tissue sections of trigeminal ganglia (TG, top panels) and cornea (bottom panels) harvested from WT (left) and CCL28^(−/−)^ deficient (right) mice following HSV-1 infection. Black arrows highlight increased epithelial disruption and inflammation within the TG and cornea.

Our results suggest that: (*i*) the CCL28/CCR10 chemokine axis may regulate the recruitment, mobilization, and functional activity of CCR10-expressing B cells and CD8+ effector memory (T_EM_) cells at sites of ocular herpes infection, including the cornea and trigeminal ganglia (TG); and (*ii*) CCL28 may play an important role in promoting the recruitment of protective CCR10^+^ memory CD8^+^ T_EM_ cells and CCR10^+^ B cells to infected ocular tissues. The accumulation of these CCR10-expressing immune cell populations at the site of HSV-1 infection may contribute to local antiviral immunity and limit viral replication, inflammation, and tissue damage during ocular herpes. These findings support a potential role for the CCL28/CCR10 axis in coordinating local memory immune responses and protecting against HSV-1-induced ocular disease.

## DISCUSSION

Recurrent ocular herpes simplex virus type 1 (HSV-1) infection remains an important cause of infectious keratitis and vision-threatening corneal disease (2, 7). Following infection of the ocular surface, HSV-1 replicates in corneal epithelial cells and subsequently establishes lifelong latency in sensory neurons of the trigeminal ganglia (TG) (7, 27). Periodic viral reactivation and anterograde transport of virus to the cornea can result in recurrent epithelial and stromal keratitis and progressive corneal damage (12, 28). Although antiviral therapy can reduce viral replication and clinical disease, it does not eliminate latent virus or provide durable protection against recurrent infection (29, 30). Therefore, the immune mechanisms that provide effective local control of HSV-1 at the ocular surface is important for the development of improved prophylactic and therapeutic vaccines.

Chemokines comprise a class of small secreted signaling proteins whose primary function is directing targeted immune cell migration during both baseline physiological state and inflammatory events (21, 31). While inflammatory chemokines govern acute immune reactivity, homeostatic variants coordinate T-cell-mediated immune responses and disease pathogenesis (21, 32). Specifically, they orchestrate the recruitment and trafficking of distinct CD8^+^ T-cell subpopulations across lymphoid structures and peripheral site lesions (33). However, significant functional redundancy exists within the chemokine network; thus, individual chemokines do not always confer protection against disease (34).

In the present study, we showed that HSV-1 infection upregulates several homeostatic mucosal chemokines CCL28 and CXCL17, in primary human corneal epithelial cells in addition to suppressing CXCL14. We investigated the role of the mucosal chemokine CCL28 and its receptor CCR10 in regulating antiviral immune responses following ocular HSV-1 infection. CCL28 is expressed by epithelial tissues and has been implicated in the recruitment of CCR10-expressing immune cells to mucosal sites (35). Based on this established function, we hypothesized that CCL28 may contribute to the localization of HSV-1-specific memory lymphocytes within ocular tissues following infection and vaccination. Our findings support this hypothesis and demonstrate an association between CCL28 expression and the accumulation of CCR10-expressing antiviral immune cells at the ocular site of HSV-1 infection.

Following ocular HSV-1 infection, we observed increased expression of CCL28 in the infected ocular tissues. This increase was accompanied by the accumulation of CCR10-expressing CD8^+^ T cells and B-cell populations. The association between increased local CCL28 expression and accumulation of CCR10^+^ lymphocytes suggests that the CCL28/CCR10 axis may provide a mechanism for directing antigen-experienced immune cells toward the ocular site of infection. This is particularly relevant to HSV-1 because the cornea represents the primary site of viral replication and pathology, while the TG serves as the principal site of viral latency (36, 37). Thus, efficient recruitment of antiviral lymphocytes to the ocular surface could provide an important first line of adaptive immune surveillance during primary infection and subsequent viral reactivation (38, 39).

In this study, we investigated the role of the mucosal chemokine CCL28 and its receptor CCR10 in regulating antiviral immunity following ocular HSV-1 infection. CCL28 (also known as mucosal-associated epithelial chemokine, MEC) is constitutively expressed by epithelial cells at multiple mucosal surfaces and signals through CCR10 and CCR3 to direct the homing of lymphocyte subsets to mucosal tissue (25, 35, 40). We hypothesized that CCL28 similarly contributes to the local positioning of HSV-1-specific memory lymphocytes at the ocular surface. Our findings support this hypothesis; HSV-1 infection increased CCL28 expression at the ocular surface, and this increase coincided with accumulation of CCR10^+^ effector-memory CD8^+^ T cells and CCR10^+^ B cells, consistent with our prior work implicating CCL28/CCR10 in effector-memory CD8^+^ T-cell recruitment to another mucosal site, the vaginal mucosa, during genital HSV-2 infection (24). Our group has also reported that CXCL17 signals through GPR35/CXCR8 to mediate CD8^+^ effector-memory and tissue-resident memory T-cell mobilization in the vaginal mucosa during genital herpes (41), and that CXCL14 similarly drives mobilization of effector-memory and tissue-resident memory T cells at this mucosal site to limit genital HSV-2 disease (42). More recently, we found that CCL25/CCR9 promotes T_RM_ accumulation and protection against ocular HSV-1 infection, and that CCL25 deficiency showed reduced expression of CCL28, CXCL14, and CXCL17 in ocular tissue (4), suggesting that these mucosal chemokines may operate within a coordinated network rather than independently. The presence of increased frequencies of HSV-1-specific CCR10^+^ memory CD8^+^ T cells in protected animals therefore suggests that CCL28-mediated recruitment may contribute to the establishment or maintenance of local antiviral immune surveillance.

The contribution of CCL28 to this response was further supported by our analysis of CCL28^(−/−)^ deficient mice, which showed reduced accumulation of CCR10+ antiviral lymphocyte populations at the ocular surface, diminished effector function (IFN-γ, TNF-α, and Granzyme B production), and increased susceptibility to primary infection and UV-B-induced reactivation. Because chemokine networks are highly redundant, the absence of CCL28 did not abolish immune-cell recruitment altogether. The reduction in specific CCR10+ memory populations suggests that CCL28 may represent one component of a broader chemokine network controlling antiviral lymphocyte trafficking to the ocular surface (43).

The reduction in CCR10^+^ B cells in CCL28^(−/−)^ deficient mice was also prominent. CCL28/CCR10 signaling has an established role in homing IgA-secreting plasma cells to mucosal epithelium (25, 44), including the gut and lactating mammary gland, and low-dose CCL28 has been shown to act as a mucosal adjuvant that boosts antigen-specific IgG and vaginal IgA when co-administered with HSV-2 glycoprotein immunogens (45, 46). Our data displayed the possibility that CCL28 coordinates the humoral arm of ocular immunity, and recruitment of CCR10^+^ B cells to the ocular mucosa could provide a local antibody source during reactivation. Future studies investigating HSV-1-specific antibody-secreting cells and local IgA and IgG responses will be important to determine whether CCL28 directly influences local humoral immunity in the cornea.

The relationship between CCL28 and tissue-resident memory T cells (T_RM_) is highly relevant to recurrent ocular herpes (4, 24, 47). T_RM_ cells provide long-term immune surveillance within peripheral tissues and can respond rapidly to local antigen exposure (48, 49). In HSV-1 infection, tissue-resident antiviral T cells have been shown to contribute to protection against recurrent infection by responding rapidly at sites of viral reactivation. T_RM_ cells have been directly implicated in controlling HSV-2 reactivation at genital epithelial and dermal sites in both mice and humans (50, 51). Our observation in ocular herpes infection also revealed that CCL28 deficiency displayed reduced CCR10^+^CD69^+^CD103^+^ CD4^+^ and CD8^+^ T-cell frequencies in ocular tissue. This suggests that the CCL28 chemokine pathway may contribute not only to the initial recruitment of circulating memory cells but also to the establishment and maintenance of a local T_RM_ compartment at the ocular surface (24, 35).

An important feature of ocular HSV-1 immunity is the need to balance antiviral protection with tissue preservation (52). Unlike many peripheral tissues, the cornea must maintain transparency for normal vision. Excessive accumulation of inflammatory cells can contribute to stromal inflammation, tissue destruction, and corneal scarring (14, 53). Therefore, simply increasing the magnitude of immune-cell infiltration may not necessarily improve disease outcome. The protective phenotype associated with CCL28 expression in our study may reflect more selective recruitment of antigen-specific memory lymphocytes rather than indiscriminate inflammation. This distinction will be important for future studies aimed at therapeutically manipulating the CCL28/CCR10 pathway.

The ability to enhance local antiviral immunity has important implications for HSV-1 vaccine development (10, 54). Previous HSV vaccine strategies have largely focused on generating systemic antibody and T-cell responses (55, 56), yet systemic immunity does not necessarily translate into efficient protection at the site of viral entry or reactivation (11, 57). The adjuvanted glycoprotein D subunit HSV vaccine candidate reached late-stage clinical efficacy, elicited strong HSV-specific neutralizing antibody responses but failed to protect against HSV-2 in a large, randomized trial, and showed only partial, HSV-1-restricted efficacy (58–60). Therefore, a successful ocular herpes vaccine may need to generate both systemic immune memory and a durable population of antiviral memory cells capable of rapidly responding within ocular tissues (61). Our findings suggest that modulation of the CCL28/CCR10 pathway could potentially improve the localization of vaccine-induced HSV-1-specific memory lymphocytes to the ocular surface. Thus, CCL28 may function not simply as an inflammatory mediator but as a component of the tissue homing that determines where protective memory cells are positioned following vaccination.

Our findings also have potential relevance to therapeutic vaccination. Because HSV-1 establishes latency in the TG, vaccination after infection is unlikely to eliminate the latent viral reservoir. However, enhancing local immune surveillance could potentially reduce viral replication following reactivation and limit the subsequent spread of virus to the cornea (12). A vaccine or immunotherapeutic strategy capable of increasing HSV-1-specific memory T-cell responses in ocular tissues may therefore reduce the severity of recurrent ocular disease even in individuals who are already infected (61). This possibility implicates further investigation using established models of HSV-1 latency and reactivation.

Several limitations should be considered. Because CCL28 signals through both CCR3 and CCR10 (26), the increased CCL28 expression and accumulation of CCR10^+^ lymphocytes demonstrate a strong association, but this limitation does not establish that CCL28 act exclusively through CCR10 to mediate protection (25). CCL28 can interact with more than one receptor, and other chemokine pathways may compensate for the loss of CCL28 (35). Then, the reduction of CCR10^+^ memory cells in CCL28^(−/−)^ deficient mice does not distinguish between impaired recruitment, altered survival, defective retention, or reduced local antigen load (44, 62). Also, phenotypic identification of memory and T_RM_ populations should be complemented by functional assays to establish their antiviral activity (63). Because ocular inflammation itself can influence chemokine expression, it will be important to determine whether CCL28 is induced directly by HSV-1 infection, by inflammatory cytokines, or as a vaccine-induced tissue-remodeling response (64, 65). Also, the route of therapeutic delivery comprising the corneal surface for translating a CCL28-based therapy offers potential advantages. As the ocular surface is directly accessible to topical and subconjunctival administration, this could allow local, sustained CCL28 delivery without systemic exposure (66, 67). Future studies should evaluate recruitment and retention of HSV-1-specific memory lymphocytes following UV-B-induced reactivation in the cornea via a topical or subconjunctival CCL28 delivery.

CCL28 may be considered as a biomarker of local antiviral immune competence (68). CCL28 is secreted at the epithelial ocular surface (69). Non-invasively sampling CCL28 levels in patients with a history of recurrent HSV-1 keratitis offers a potential correlate of local immune surveillance capacity or recurrence risk (7). The translation of our current findings, analogous to the use of local cytokine/chemokine profiling in other recurrent mucosal viral infections, may offer a diagnostic tool (24, 68). CCL28 also demonstrates a direct antimicrobial role against Gram-positive and Gram-negative bacteria and fungi, contributing to innate epithelial defense at mucosal surfaces (35, 70–72). HSV keratitis also conveys a recognized risk of secondary bacterial infection, due to epithelial breakdown (28). CCL28 deficiency may increase the disease risk through loss of antimicrobial function and impaired lymphocyte recruitment. This dual role was not distinguished in the present study and should be studied when interpreting the increased ocular pathology observed in CCL28^(−/−)^ deficient mice. Furthermore, prior research showed a pivotal role of estrogen in conferring host defense against genital tract infections by modulating mucosal epithelial cell (MEC) expression of CCL28 within the uterus (73, 74). Consequently, investigating how sex steroids such as estrogen influence CCL28 activity represents a promising future study. The clinical role of CCL28 has also been reported in other pathological conditions, including the maintenance of cardiac function following myocardial infarction and protection against ischemia–reperfusion injury in the kidney (75). Moreover, CCL28-CCR10 signaling has been implicated in NF-κB pathway-regulated neuroprotection following spinal cord injury, potentially through the recruitment and immunosuppressive activity of regulatory T (Treg) cells (76). Similarly, we recently reported NF-κB/OCT-1 mediated upregulation of CXCL14 in genital herpes (42). These findings suggest that similar CCL28–CCR10-mediated immunoregulatory mechanisms may also contribute to tissue protection and immune regulation during ocular herpes and require further investigation.

In summary, our findings identify the CCL28/CCR10 chemokine axis as an important component of local antiviral immunity following ocular HSV-1 infection. Increased CCL28 expression was associated with enhanced accumulation of HSV-1-specific CCR10-expressing memory T cells and B-cell populations at the ocular site of infection, whereas loss of CCL28 resulted in impaired localization of these immune populations and compromised antiviral protection. These observations suggest that CCL28 contributes to the establishment of a localized antiviral immune environment in the ocular mucosa by promoting the recruitment and retention of antigen-experienced lymphocytes. These results provide a rationale for further defining how CCL28 induces ocular T_RM_ formation, local humoral immunity, and reactivation control may provide new opportunities to develop mucosal chemokine-based vaccines and immunotherapeutic approaches capable of generating durable and protective immunity against recurrent ocular herpes.

## MATERIALS AND METHODS

### Virus propagation and titration

Vero cells were maintained in Dulbecco’s Modified Eagle Medium (DMEM) supplemented with 10% fetal bovine serum (FBS) and 1% penicillin-streptomycin and were used for propagation and quantification of HSV-1. The HSV-1 strain McKrae was propagated in Vero cells as described previously(77, 78). Briefly, virus stocks were prepared from infected cell cultures and stored at −80°C. Viral titers were determined by plaque assay on Vero cell monolayers and expressed as plaque-forming units (PFU)/ml. The same virus stock was used for all experimental infections to minimize variability between experiments. HSV-1 was selected for these studies because it is the predominantly associate with ocular infection and can establish latency in the trigeminal ganglia following corneal infection.

### HSV-1 infection of human primary corneal epithelial cells

Primary human corneal epithelial cells (HCECs; ATCC® PCS-700-010™) were obtained from the American Type Culture Collection (ATCC, Manassas, VA, USA) and maintained according to the manufacturer’s recommendations. Cells were cultured in the recommended corneal epithelial cell growth medium and maintained under standard cell culture conditions. The corneal epithelial cells were plated in 48-well plates at a density of 1 × 10⁴ cells per well in 300 μl of growth medium and maintained until reaching approximately 80– 90% confluence. The cells were then infected with HSV-1 McKrae at the multiplicity of infection of 2.5 in 100 μl of growth medium for 1 h. Following viral adsorption, the cells were washed four times with PBS to remove unbound virus. HSV-1 induced chemokine expression was evaluated using quantitative real-time PCR (qRT-PCR) as described earlier (42). Post-infection, total RNA was isolated from the cells collected at various time-point, reverse-transcribed into cDNA, and analyzed using gene-specific forward and reverse primers. CCL25: 5’ TTATTCGTCCAGGTGCCCAG 3’ and 5’ GGCAGCAGTCCTCAAAGACA 3’, CCL28: 5’ CTTGTGTTGCTGTCAGTGCC 3’ and 5’ GGCTTCTGA GGCATGTAGGG 3’, CXCL17- 5’ CAGTCTTAGCCTGTGCCCTC 3’ and 5’ GGCAGACCCCATTTGAAGGA 3’, CXCL14- 5’ CTAAGATGACCATGCGCCCT 3’ and 5’ AATGCGGCATATACTGGGGG 3’, GAPDH-5’ GGTGTGAACCATGAGAAGTATGAC A 3’ and 5’ GGTGCAGGAGGCATTGCT 3’.

### Mice

C57BL/6 (B6) wild-type mice, 6–8 weeks of age, were obtained from the Jackson Laboratory (Bar Harbor, ME, USA). CCL28^(−/−)^ deficient mice were obtained from Dr. Takashi Nakayama, Kindai University, Japan, and bred at the University of California, Irvine (UCI) animal facility. CCL28^(−/−)^ deficient mice and age-matched wild-type mice, 6–8 weeks of age, were used for the recurrent ocular HSV-1 infection studies. Mice were housed under specific pathogen-free conditions with access to food and water ad libitum. All animal experiments were conducted in accordance with the Guide for the Care and Use of Laboratory Animals published by the National Institutes of Health and were approved by the Institutional Animal Care and Use Committee of the University of California, Irvine (UCI). All procedures were performed in accordance with the approved institutional animal-use protocol (IACUC protocol # AUP-22-086).

### Ocular HSV-1 infection and recurrent disease model

C57BL/6 wild-type (WT) and CCL28^(−/−)^ deficient mice were used to investigate the role of the CCL28/CCR10 chemokine axis during recurrent ocular HSV-1 infection. Mice were inoculated ocularly with HSV-1 (strain McKrae) following the established ocular infection protocol (4). After infection, mice were monitored daily for the development and progression of ocular disease. Clinical disease was evaluated using an established corneal scoring system based on the severity and extent of corneal opacity, inflammation, and neovascularization, as described previously (79, 80). To investigate the development of recurrent ocular herpes, mice that survived the primary infection were maintained for the indicated period following resolution of acute disease. Latent HSV-1 infection was allowed to establish in the trigeminal ganglia (TG). To investigate recurrent ocular HSV-1 disease, mice that survived primary ocular infection and established latent infection were subjected to UVB treatment to induce viral reactivation, as described previously(81). Briefly, mice were treated with UVB on day 21, the eyes of the mice were exposed to UVB light at a dose of 250 mJ/cm² for 60 seconds on the transilluminator. Following UVB exposure, animals were monitored daily for clinical evidence of recurrent ocular disease, including corneal opacity, inflammation, and neovascularization. Ocular disease was scored using the established clinical scoring system described above. To determine the effect of CCL28 deficiency on recurrent ocular HSV-1 disease, age-matched WT and CCL28^(−/−)^ deficient mice that survived primary infection were subjected to UVB-induced reactivation. Ocular swabs were collected at the indicated time points following UVB treatment to assess recurrent viral shedding. Viral burden in ocular tissues and trigeminal ganglia (TG) was determined by quantitative PCR. The UVB-induced HSV-1 reactivated mice were euthanized at the end time point on day 41, and corneas/ocular tissues, TG, and spleens were collected for virological and immunological analyses.

### Monitoring of ocular herpes infection and disease scoring in mice

Mice that survived primary infection and established latency were monitored for disease progression following the induction of HSV-1 reactivation. Viral shedding was assessed by collecting ocular surface swabs from infected mice post UVB reactivation on day 21. Briefly, the ocular surface was gently swabbed using sterile swabs, and samples were stored at −80°C until viral titration. Infectious HSV-1 was quantified by quantitative viral titration. Mice were also monitored for clinical signs of ocular disease following HSV-1 infection. Corneal disease was evaluated by slit-lamp examination and scored based on the extent of corneal opacity and inflammation using a scale of 0–4: 0, no detectable disease; 1, mild corneal opacity with iris details clearly visible; 2, moderate opacity with partial obscuring of iris details; 3, severe corneal opacity with iris details largely obscured; and 4, complete corneal opacity and/or corneal perforation. Clinical observations also included corneal edema, epithelial lesions, stromal inflammation, and neovascularization. Mice were additionally evaluated for signs of neurological disease, including abnormal posture, reduced mobility, urinary or fecal retention, hind-limb weakness, paresis, or paralysis. Neurological disease was scored on a scale ranging from 0 (no detectable neurological abnormalities) to 4 (severe neurological impairment or hind-limb paralysis). Animals reaching the predetermined humane endpoint or exhibiting severe disease were euthanized in accordance with the approved animal protocol.

### Quantitative real-time PCR for HSV-1 DNA quantification

Genomic DNA was extracted from corneal swab samples using *Quick*-DNA Viral Kit (Zymo Research, Irvine, CA) according to the manufacturer’s instructions. DNA concentration and purity were determined before quantitative PCR (qPCR). HSV-1 DNA was quantified using a SYBR Green-based real-time PCR assay targeting a conserved HSV-1 genomic region. The qPCR reaction was performed using SYBR Green master mix containing DNA polymerase, dNTPs, MgCl₂, and SYBR Green fluorescent dye, together with HSV-1-specific forward and reverse primers. Each reaction contained the SYBR Green master mix, HSV-1-specific primers, and the appropriate amount of template DNA in a final reaction volume of 10 µl. The primer sequences used for HSV-1 detection were: Forward Primer: 5’-CATCACCGACCCGGAGAGGGAC-3’ and Reverse Primer: 5’-GGGCCAGGCGCTTGTTGGTGTA-3’. A no-template control was included in each assay to monitor potential contamination. Real-time PCR was performed using an Applied Biosystems QuantStudio 5 quantitative PCR system with the following cycling conditions: initial denaturation at 95 °C for 20 sec, followed by 40 cycles of denaturation at 95 °C for 1 s, primer annealing at 60 °C for 20 s, and extension at 95 °C for 1 s. A melt-curve analysis was performed at the end of each run to confirm the specificity of the amplified product. HSV-1 DNA levels were quantified using a standard curve generated from serial dilutions of a known HSV-1 DNA standard.

### Flow cytometry

Single-cell suspensions were prepared from the trigeminal ganglia (TG), corneas, and spleens of HSV-1-infected mice for flow cytometric analysis. Briefly, TG and corneal tissues were collected at the indicated time points and processed to obtain single-cell suspensions using enzymatic digestion with Collagenase D (Millipore Sigma, St. Louis, MO), under conditions optimized for each tissue. Spleens were mechanically dissociated to generate single-cell suspensions, followed by removal of erythrocytes using ACK red blood cell lysis buffer (Gibco, Waltham, MA). Cell suspensions were passed through cell strainers, washed with PBS, and resuspended in FACS buffer.

For surface staining, approximately 1 × 10⁶ cells were incubated with fluorochrome-conjugated monoclonal antibodies against the indicated cell-surface markers in PBS containing 1% FBS and 0.1% sodium azide (FACS buffer) for 45 min at 4°C in the dark. The antibody panel included markers for T-cell and B-cell identification and characterization, including anti-mouse CD45 (clone 30-F11), CD3 (clone 17A-2), CD4, CD8, CD44, CD62L, CD69, CD103, and B220 (BD Biosciences). Following staining, cells were washed twice with FACS buffer and subsequently fixed in PBS containing 2 % paraformaldehyde (Sigma-Aldrich, St. Louis, MO). For intracellular staining, cells were treated with Cytofix/Cytoperm solution (BD Biosciences, USA) following surface staining and incubated for 45 min at 4 °C with antibodies against IFN-γ, TNF-α, and Granzyme B. Cells were then washed with FACS buffer and fixed again with 2% paraformaldehyde. A total of 100,000 lymphocyte-gated PBMC events were acquired on a BD Fortessa X20 flow cytometer (Becton Dickinson, Mountain View, CA). Data were analyzed using FlowJo software version 10.10.0 (Becton Dickinson, Ashland, OR). Appropriate fluorescence-minus-one (FMO) and unstained controls were included for gating and determination of marker-positive populations.

### Histopathology and immunohistochemistry of trigeminal ganglia and corneal tissues

Trigeminal ganglia (TG) and corneal tissues were collected from mice on day 41 following HSV-1 infection and processed for histopathological and immunohistochemical analyses. For hematoxylin and eosin (H&E) staining, tissue sections were deparaffinized and rehydrated, followed by staining with hematoxylin and eosin according to standard histological procedures. H&E-stained sections were examined for tissue architecture, inflammatory cell infiltration, epithelial disruption, stromal inflammation, and other pathological changes associated with HSV-1 infection. For immunohistochemical analysis, tissue sections were deparaffinized and rehydrated, followed by antigen retrieval and blocking of nonspecific binding. Sections were incubated overnight at 4°C with primary antibodies specific for mouse CCL28, CCR10, CD4, and CD8. After washing with PBS, sections were incubated with Alexa fluor-conjugated secondary antibodies for 1 hour in the dark. ProLong™ Diamond Antifade Mountant was used for mounting slides. For hematoxylin and eosin (H&E) staining, tissues were fixed in 4% paraformaldehyde (PFA) for 48 h and subsequently transferred to 70% ethanol. The tissues were processed, embedded in paraffin, and sectioned at a thickness of 8 μm. Sections were deparaffinized and rehydrated, followed by H&E staining for histopathological evaluation. Images were captured on a Keyence BZ-X810 fluorescence microscope.

### Statistical analysis

Data for each assay were compared by ANOVA and Student’s *t*-test using GraphPad Prism version 5 (La Jolla, CA). As previously described, differences between the groups were identified by ANOVA and multiple comparison procedures (33, 34). Data are expressed as the mean ± SD. Results were considered statistically significant at a *P* value of ≤ 0.05.

## ACKNOWLEDGEMENTS

This work is supported by Public Health Service Research R01 Grants EY026103, EY019896, and EY024618 from the National Eye Institute (NEI), AI158060, AI150091, AI143348, AI147499, AI143326, AI138764, AI124911, and AI110902 from the National Institutes of Allergy and Infectious Diseases (NIAID) to LBM, and in part by The Discovery Center for Eye Research (DCER) and the Research to Prevent Blindness (RPB) grant. This work is dedicated to the memory of the late Professor Steven L. Wechsler “Steve” (1948-2016), whose numerous pioneering works on herpes infection and immunity laid the foundation for this line of research. We thank the NIH Tetramer Facility (Emory University, Atlanta, GA) for providing the Tetramers used in this study.

---

**Supplementary Figure 1. Example of a flow cytometry gating strategy and phenotypic analysis of T cell and B cell subsets:** (**A**) Representative flow cytometry plots illustrating the gating hierarchy to identify lymphocyte subsets. Cells were first gated on forward scatter (FSC-A) vs. side scatter (SSC-A) to isolate the lymphocyte population, followed by singlet discrimination (FSC-H vs. FSC-A). CD45+ leukocytes were subsequently gated to identify B cells (B220^+^) and T lymphocytes (CD3^+^), with further resolution of T cells into CD4^+^ and CD8^+^ T cell populations. (**B**) Assessment of homing receptor expression showing frequencies of B220^+^, CD3^+^, CD4^+^, and CD8^+^ expressing CCR10 (%). (**C**) Representative flow plots displaying activation and tissue-resident memory (T_RM_) marker expression and naive, central memory (T_CM_), and effector memory (T_EM_)-CD69, CD103, CD62L, CD44 gated on CD4^+^ and CD8^+^ T cells. Intracellular cytokine and cytotoxic molecule staining of CD4^+^ and CD8^+^ T cells evaluating expression of Granzyme B (GzmB), Interferon-gamma (IFN-γ), and Tumor Necrosis Factor (TNF-*a*).

## REFERENCES

1. Lekbach Y, Chentoufi AA, Prakash S, Karan S, Quadiri A, Hormi-Carver KK, Dorotta JC, BenMohamed L. 2026. Role of mucosal chemokines in the development of tissue-resident CD4(+) and CD8(+) T(RM) cells to fend off herpes simplex infections. Front Immunol 17:1661640.

2. McCormick I, James C, Welton NJ, Mayaud P, Turner KME, Gottlieb SL, Foster A, Looker KJ. 2022. INCIDENCE OF HERPES SIMPLEX VIRUS KERATITIS AND OTHER OCULAR DISEASE: GLOBAL REVIEW AND ESTIMATES. Ophthalmic Epidemiol 29:353–362.

3. Quadiri A, Lekbach Y, Elfatimi E, Prakash S, Vahed H, Karan S, Rehman A, Ng SXL, Maurya C, Chow R, BenMohamed L. 2025. The Path Towards Effective Long-Lasting Tissue-Targeted Prime/Pull/Keep Herpes Simplex Therapeutic Vaccines. Vaccines (Basel) 13.

4. Rahman A, Prakash S, Karan S, Ng SXL, Park G, Maurya C, Garcia A, Intharachalit K, Hwang J, Tang CTY, Chow RA, Suoth BS, Liao EJ, BenMohamed L. 2026. CCL25 chemokine promotes antiviral tissue resident CD4(+) and CD8(+) effector memory T(RM) cells associated with a reduction of ocular herpes infection and disease: a potential gut-eye axis in herpes immunity. Front Immunol 17:1872553.

5. Rajasagi NK, Rouse BT. 2018. Application of our understanding of pathogenesis of herpetic stromal keratitis for novel therapy. Microbes Infect 20:526–530.

6. Musa M, Enaholo E, Aluyi-Osa G, Atuanya GN, Spadea L, Salati C, Zeppieri M. 2024. Herpes simplex keratitis: A brief clinical overview. World J Virol 13:89934.

7. Farooq AV, Shukla D. 2012. Herpes simplex epithelial and stromal keratitis: an epidemiologic update. Surv Ophthalmol 57:448–62.

8. Al-Dujaili LJ, Clerkin PP, Clement C, McFerrin HE, Bhattacharjee PS, Varnell ED, Kaufman HE, Hill JM. 2011. Ocular herpes simplex virus: how are latency, reactivation, recurrent disease and therapy interrelated? Future Microbiol 6:877–907.

9. Held K, Derfuss T. 2011. Control of HSV-1 latency in human trigeminal ganglia--current overview. J Neurovirol 17:518–27.

10. Su D, Han L, Shi C, Li Y, Qian S, Feng Z, Yu L. 2024. An updated review of HSV-1 infection-associated diseases and treatment, vaccine development, and vector therapy application. Virulence 15:2425744.

11. Bai L, Xu J, Zeng L, Zhang L, Zhou F. 2024. A review of HSV pathogenesis, vaccine development, and advanced applications. Molecular Biomedicine 5:35.

12. Greenan E, Gallagher S, Khalil R, Murphy CC, J NG-D. 2021. Advancing Our Understanding of Corneal Herpes Simplex Virus-1 Immune Evasion Mechanisms and Future Therapeutics. Viruses 13.

13. Abdelfattah NS, Amgad M, Zayed AA. 2016. Host immune cellular reactions in corneal neovascularization. Int J Ophthalmol 9:625–33.

14. Fortingo N, Melnyk S, Sutton SH, Watsky MA, Bollag WB. 2022. Innate Immune System Activation, Inflammation and Corneal Wound Healing. Int J Mol Sci 23.

15. Osborne L, Dunai C, Huang Y, Egbe FN, Turtle L, Michael BD, Ellul MA. 2026. The cell-mediated adaptive immune response to herpes simplex virus type 1 encephalitis: mechanisms and clinical implications. Front Immunol 17:1751672.

16. Zhang J, Liu H, Wei B. 2017. Immune response of T cells during herpes simplex virus type 1 (HSV-1) infection. J Zhejiang Univ Sci B 18:277–288.

17. Wang L, Wang R, Xu C, Zhou H. 2020. Pathogenesis of Herpes Stromal Keratitis: Immune Inflammatory Response Mediated by Inflammatory Regulators. Front Immunol 11:766.

18. Chentoufi AA, BenMohamed L, Van De Perre P, Ashkar AA. 2012. Immunity to ocular and genital herpes simplex viruses infections. Clin Dev Immunol 2012:732546.

19. Khanna KM, Bonneau RH, Kinchington PR, Hendricks RL. 2003. Herpes simplex virus-specific memory CD8+ T cells are selectively activated and retained in latently infected sensory ganglia. Immunity 18:593–603.

20. Lam JH, Smith FL, Baumgarth N. 2020. B Cell Activation and Response Regulation During Viral Infections. Viral Immunol 33:294–306.

21. Sokol CL, Luster AD. 2015. The chemokine system in innate immunity. Cold Spring Harb Perspect Biol 7.

22. Hernandez-Ruiz M, Zlotnik A. 2017. Mucosal Chemokines. J Interferon Cytokine Res 37:62–70.

23. Smith JB, Herbert JJ, Truong NR, Cunningham AL. 2022. Cytokines and chemokines: The vital role they play in herpes simplex virus mucosal immunology. Frontiers in Immunology Volume 13–2022.

24. Dhanushkodi NR, Prakash S, Quadiri A, Zayou L, Srivastava R, Tran J, Dang V, Shaik AM, Chilukurri A, Suzer B, Vera P, Sun M, Nguyen P, Lee A, Salem A, Loi J, Singer M, Nakayama T, Vahed H, Nesburn AB, BenMohamed L. 2023. Mucosal CCL28 Chemokine Improves Protection against Genital Herpes through Mobilization of Antiviral Effector Memory CCR10+CD44+ CD62L-CD8+ T Cells and Memory CCR10+B220+CD27+ B Cells into the Infected Vaginal Mucosa. J Immunol 211:118–129.

25. Chen Z, Kim SJ, Essani AB, Volin MV, Vila OM, Swedler W, Arami S, Volkov S, Sardin LV, Sweiss N, Shahrara S. 2015. Characterising the expression and function of CCL28 and its corresponding receptor, CCR10, in RA pathogenesis. Ann Rheum Dis 74:1898–906.

26. Mehta MK, Gupta S, Fatima T, Selvam R, Chandra S, Singh D, Sivakumar N. 2024. An Increase in the Chemokine Mediators (CCL28 and CCR10) Associated with the Progression of Oral Squamous Cell Carcinoma: A Cross-Sectional Investigation. Indian J Otolaryngol Head Neck Surg 76:5717–5724.

27. Chentoufi AA, Kritzer E, Tran MV, Dasgupta G, Lim CH, Yu DC, Afifi RE, Jiang X, Carpenter D, Osorio N, Hsiang C, Nesburn AB, Wechsler SL, BenMohamed L. 2011. The herpes simplex virus 1 latency-associated transcript promotes functional exhaustion of virus-specific CD8+ T cells in latently infected trigeminal ganglia: a novel immune evasion mechanism. J Virol 85:9127–38.

28. Lobo AM, Agelidis AM, Shukla D. 2019. Pathogenesis of herpes simplex keratitis: The host cell response and ocular surface sequelae to infection and inflammation. Ocul Surf 17:40–49.

29. Du S, Hu X, Li P, Xu S, Kim M, Liu X, Zhan P. 2026. Antiviral drug discovery and development: challenges and future directions. Signal Transduct Target Ther 11.

30. Li Z, Zhang Y, Zheng Y, Wang H, Xu C, He Q. 2026. Therapeutic Vaccines for Chronic Viral Infections: From Immune Modulation to Clinical Translation. Vaccines 14:507.

31. Hughes CE, Nibbs RJB. 2018. A guide to chemokines and their receptors. Febs j 285:2944–2971.

32. Chen K, Bao Z, Tang P, Gong W, Yoshimura T, Wang JM. 2018. Chemokines in homeostasis and diseases. Cell Mol Immunol 15:324–334.

33. Farsakoglu Y, McDonald B, Kaech SM. 2021. Motility Matters: How CD8(+) T-Cell Trafficking Influences Effector and Memory Cell Differentiation. Cold Spring Harb Perspect Biol 13.

34. Dyer DP. 2020. Understanding the mechanisms that facilitate specificity, not redundancy, of chemokine-mediated leukocyte recruitment. Immunology 160:336–344.

35. Mohan T, Deng L, Wang BZ. 2017. CCL28 chemokine: An anchoring point bridging innate and adaptive immunity. Int Immunopharmacol 51:165–170.

36. Kennedy DP, Clement C, Arceneaux RL, Bhattacharjee PS, Huq TS, Hill JM. 2011. Ocular herpes simplex virus type 1: is the cornea a reservoir for viral latency or a fast pit stop? Cornea 30:251–9.

37. Farooq AV, Shukla D. 2011. Corneal latency and transmission of herpes simplex virus-1. Future Virol 6:101–108.

38. Petrillo F, Petrillo A, Sasso FP, Schettino A, Maione A, Galdiero M. 2022. Viral Infection and Antiviral Treatments in Ocular Pathologies. Microorganisms 10.

39. de Paiva CS, St. Leger AJ, Caspi RR. 2022. Mucosal immunology of the ocular surface. Mucosal Immunology 15:1143–1157.

40. Meurens F, Whale J, Brownlie R, Dybvig T, Thompson DR, Gerdts V. 2007. Expression of mucosal chemokines TECK/CCL25 and MEC/CCL28 during fetal development of the ovine mucosal immune system. Immunology 120:544–55.

41. Srivastava R, Hernández-Ruiz M, Khan AA, Fouladi MA, Kim GJ, Ly VT, Yamada T, Lam C, Sarain SAB, Boldbaatar U, Zlotnik A, Bahraoui E, BenMohamed L. 2018. CXCL17 Chemokine-Dependent Mobilization of CXCR8(+)CD8(+) Effector Memory and Tissue-Resident Memory T Cells in the Vaginal Mucosa Is Associated with Protection against Genital Herpes. J Immunol 200:2915–2926.

42. Lekbach Y, Prakash S, Vahed H, Quadiri A, Rahman A, El Fatimi EH, Dorotta JC, Suoth BS, Maurya C, Garcia A, Park G, BenMohamed L. 2026. NF-κB/OCT-1 Mediated Upregulation of CXCL14 Chemokine Mobilizes Mucosal Effector Memory CD44+CD62L-CD4+ and CD8+ TEM Cells, and NK Cells Associated with Protection Against Genital Herpes. Pathog Immun 11:191–222.

43. Enriquez-De-Salamanca A, Domínguez-López A, Blanco-Vázquez M, Calderon-Garcia AA, Garcia-Vazquez C, González-García MJ, Calonge M. 2023. Analysis of the mucosal chemokines CCL28, CXCL14, and CXCL17 in dry eye. Investigative Ophthalmology & Visual Science 64:685–685.

44. Hu S, Yang K, Yang J, Li M, Xiong N. 2011. Critical roles of chemokine receptor CCR10 in regulating memory IgA responses in intestines. Proceedings of the National Academy of Sciences 108:E1035–E1044.

45. Morteau O, Gerard C, Lu B, Ghiran S, Rits M, Fujiwara Y, Law Y, Distelhorst K, Nielsen EM, Hill ED, Kwan R, Lazarus NH, Butcher EC, Wilson E. 2008. An indispensable role for the chemokine receptor CCR10 in IgA antibody-secreting cell accumulation. J Immunol 181:6309–15.

46. Willuveit AL, Stefanini ACB, Matozo T, Marti LC. 2025. CCR10: a comprehensive review of its function, phylogeny, role in immune cell trafficking and disease pathogenesis. Frontiers in Immunology Volume 16–2025.

47. O’Neil TR, Hu K, Truong NR, Arshad S, Shacklett BL, Cunningham AL, Nasr N. 2021. The Role of Tissue Resident Memory CD4 T Cells in Herpes Simplex Viral and HIV Infection. Viruses 13:359.

48. Xie D, Lu G, Mai G, Guo Q, Xu G. 2025. Tissue-resident memory T cells in diseases and therapeutic strategies. MedComm (2020) 6:e70053.

49. Schreiner D, King CG. 2018. CD4+ Memory T Cells at Home in the Tissue: Mechanisms for Health and Disease. Front Immunol 9:2394.

50. Zhu J, Miner MD. 2024. Local Power: The Role of Tissue-Resident Immunity in Human Genital Herpes Simplex Virus Reactivation. Viruses 16.

51. Peng T, Phasouk K, Sodroski CN, Sun S, Hwangbo Y, Layton ED, Jin L, Klock A, Diem K, Magaret AS, Jing L, Laing K, Li A, Huang M-L, Mertens M, Johnston C, Jerome KR, Koelle DM, Wald A, Knipe DM, Corey L, Zhu J. 2021. Tissue-Resident-Memory CD8+ T Cells Bridge Innate Immune Responses in Neighboring Epithelial Cells to Control Human Genital Herpes. Frontiers in Immunology Volume 12–2021.

52. Ren J, Antony F, Rouse BT, Suryawanshi A. 2023. Role of Innate Interferon Responses at the Ocular Surface in Herpes Simplex Virus-1-Induced Herpetic Stromal Keratitis. Pathogens 12:437.

53. Yeung V, Boychev N, Farhat W, Ntentakis DP, Hutcheon AEK, Ross AE, Ciolino JB. 2022. Extracellular Vesicles in Corneal Fibrosis/Scarring. Int J Mol Sci 23.

54. Singer M, Husseiny MI. 2024. Immunological Considerations for the Development of an Effective Herpes Vaccine. Microorganisms 12.

55. Dropulic LK, Cohen JI. 2012. The challenge of developing a herpes simplex virus 2 vaccine. Expert Rev Vaccines 11:1429–40.

56. Chang JY, Balch C, Oh HS. 2024. Toward the Eradication of Herpes Simplex Virus: Vaccination and Beyond. Viruses 16.

57. Truong NR, Smith JB, Sandgren KJ, Cunningham AL. 2019. Mechanisms of Immune Control of Mucosal HSV Infection: A Guide to Rational Vaccine Design. Front Immunol 10:373.

58. Mahant AM, Guerguis S, Blevins TP, Cheshenko N, Gao W, Anastos K, Belshe RB, Herold BC. 2022. Failure of Herpes Simplex Virus Glycoprotein D Antibodies to Elicit Antibody-Dependent Cell-Mediated Cytotoxicity: Implications for Future Vaccines. J Infect Dis 226:1489–1498.

59. Leroux-Roels G, Dobson S, Bernstein DI, Fowler S, Romanowski B, Leroux-Roels I, Cheuvart B, Heineman T, Dubin G. 2013. Clinical evaluation to confirm the manufacturing consistency of three lots of an adjuvanted glycoprotein D genital herpes vaccine in healthy seronegative pre-teen and adolescent girls: A phase III multi-center double-blind randomized trial. Trials in Vaccinology 2:10–18.

60. Belshe RB, Leone PA, Bernstein DI, Wald A, Levin MJ, Stapleton JT, Gorfinkel I, Morrow RL, Ewell MG, Stokes-Riner A, Dubin G, Heineman TC, Schulte JM, Deal CD. 2012. Efficacy results of a trial of a herpes simplex vaccine. N Engl J Med 366:34–43.

61. Royer DJ, Hendrix JF, Larabee CM, Reagan AM, Sjoelund VH, Robertson DM, Carr DJJ. 2019. Vaccine-induced antibodies target sequestered viral antigens to prevent ocular HSV-1 pathogenesis, preserve vision, and preempt productive neuronal infection. Mucosal Immunology 12:827–839.

62. Gary EN, Kathuria N, Makurumidze G, Curatola A, Ramamurthi A, Bernui ME, Myles D, Yan J, Pankhong P, Muthumani K, Haddad E, Humeau L, Weiner DB, Kutzler MA. 2020. CCR10 expression is required for the adjuvant activity of the mucosal chemokine CCL28 when delivered in the context of an HIV-1 Env DNA vaccine. Vaccine 38:2626–2635.

63. Heeg M, Goldrath AW. 2023. Insights into phenotypic and functional CD8(+) T(RM) heterogeneity. Immunol Rev 316:8–22.

64. Wuest TR, Carr DJ. 2008. The role of chemokines during herpes simplex virus-1 infection. Front Biosci 13:4862–72.

65. Carr DJ, Tomanek L. 2006. Herpes simplex virus and the chemokines that mediate the inflammation. Curr Top Microbiol Immunol 303:47–65.

66. Shastri DH, Silva AC, Almeida H. 2023. Ocular Delivery of Therapeutic Proteins: A Review. Pharmaceutics 15.

67. Mofidfar M, Abdi B, Ahadian S, Mostafavi E, Desai TA, Abbasi F, Sun Y, Manche EE, Ta CN, Flowers CW. 2021. Drug delivery to the anterior segment of the eye: A review of current and future treatment strategies. Int J Pharm 607:120924.

68. Kaibori Y, Tamoto S, Okuda S, Matsuo K, Nakayama T, Nagakubo D. 2024. CCL28: A Promising Biomarker for Assessing Salivary Gland Functionality and Maintaining Healthy Oral Environments. Biology 13:147.

69. Domínguez-López A, Blanco-Vázquez M, Calderón-García AÁ, García-Vázquez C, González-García MJ, Calonge M, Enríquez-de-Salamanca A. 2024. Analysis of the mucosal chemokines CCL28, CXCL14, and CXCL17 in dry eye disease: An in vitro and clinical investigation. Experimental Eye Research 241:109854.

70. Liu B, Wilson E. 2010. The antimicrobial activity of CCL28 is dependent on C-terminal positively-charged amino acids. Eur J Immunol 40:186–96.

71. Berri M, Virlogeux-Payant I, Chevaleyre C, Melo S, Zanello G, Salmon H, Meurens F. 2014. CCL28 involvement in mucosal tissues protection as a chemokine and as an antibacterial peptide. Developmental & Comparative Immunology 44:286–290.

72. Walker GT, Perez-Lopez A, Silva S, Lee MH, Bjånes E, Dillon N, Brandt SL, Gerner RR, Melchior K, Norton GJ, Argueta FA, Dela Pena F, Park L, Sosa-Hernandez VA, Cervantes-Diaz R, Romero-Ramirez S, Cartelle Gestal M, Maravillas-Montero JL, Nuccio S-P, Nizet V, Raffatellu M. 2024. CCL28 modulates neutrophil responses during infection with mucosal pathogens. eLife 13:e78206.

73. Cha HR, Ko HJ, Kim ED, Chang SY, Seo SU, Cuburu N, Ryu S, Kim S, Kweon MN. 2011. Mucosa-associated epithelial chemokine/CCL28 expression in the uterus attracts CCR10+ IgA plasma cells following mucosal vaccination via estrogen control. J Immunol 187:3044–52.

74. Choi Y, Seo H, Han J, Yoo I, Kim J, Ka H. 2016. Chemokine (C-C Motif) Ligand 28 and Its Receptor CCR10: Expression and Function at the Maternal-Conceptus Interface in Pigs. Biology of Reproduction 95.

75. Yang K, Chen H, Lyu Y, Wei W, Wei X, Ling Y, Lin B, Zhou G, Chen J, Shi J, Gao R, Lin K. 2025. CCL28 contributes to angiogenesis and cardiac repair through CCR10+ endothelial cells after myocardial infarction in male mice. Nature Communications 16:9262.

76. Wang P, Qi X, Xu G, Liu J, Guo J, Li X, Ma X, Sun H. 2019. CCL28 promotes locomotor recovery after spinal cord injury via recruiting regulatory T cells. Aging 11:7402–7415.

77. Nguyen HM, Sah N, Humphrey MRM, Rabkin SD, Saha D. 2021. Growth, Purification, and Titration of Oncolytic Herpes Simplex Virus. J Vis Exp doi:10.3791/62677.

78. Fabiani M, Limongi D, Palamara AT, De Chiara G, Marcocci ME. 2017. A Novel Method to Titrate Herpes Simplex Virus-1 (HSV-1) Using Laser-Based Scanning of Near-Infrared Fluorophores Conjugated Antibodies. Frontiers in Microbiology Volume 8–2017.

79. Pepperberg IM. 1986. SENSITIVE PERIODS, SOCIAL-INTERACTION, AND SONG ACQUISITION - THE DIALECTICS OF DIALECTS. Behavioral and Brain Sciences 9:756–757.

80. Muller E, Feinberg L, Woronkowicz M, Roberts HW. 2026. Corneal Neovascularization: Pathogenesis, Current Insights and Future Strategies. Biology 15:136.

81. Yin X-T, Hartman A, Sirajuddin N, Shukla D, Leger AS, Keadle TL, Stuart PM. 2024. UVB induced reactivation leads to HSV1 in the corneas of virtually all latently infected mice and requires STING to develop corneal disease. Scientific Reports 14:6859.

