## Supplementary material for "Tissue-Resident and Effector-Memory Lymphocyte Recruitment to the Ocular Mucosa Requires CCL28/CCR10 Signaling During Recurrent HSV-1 Infection": Suplemenal Figure 1

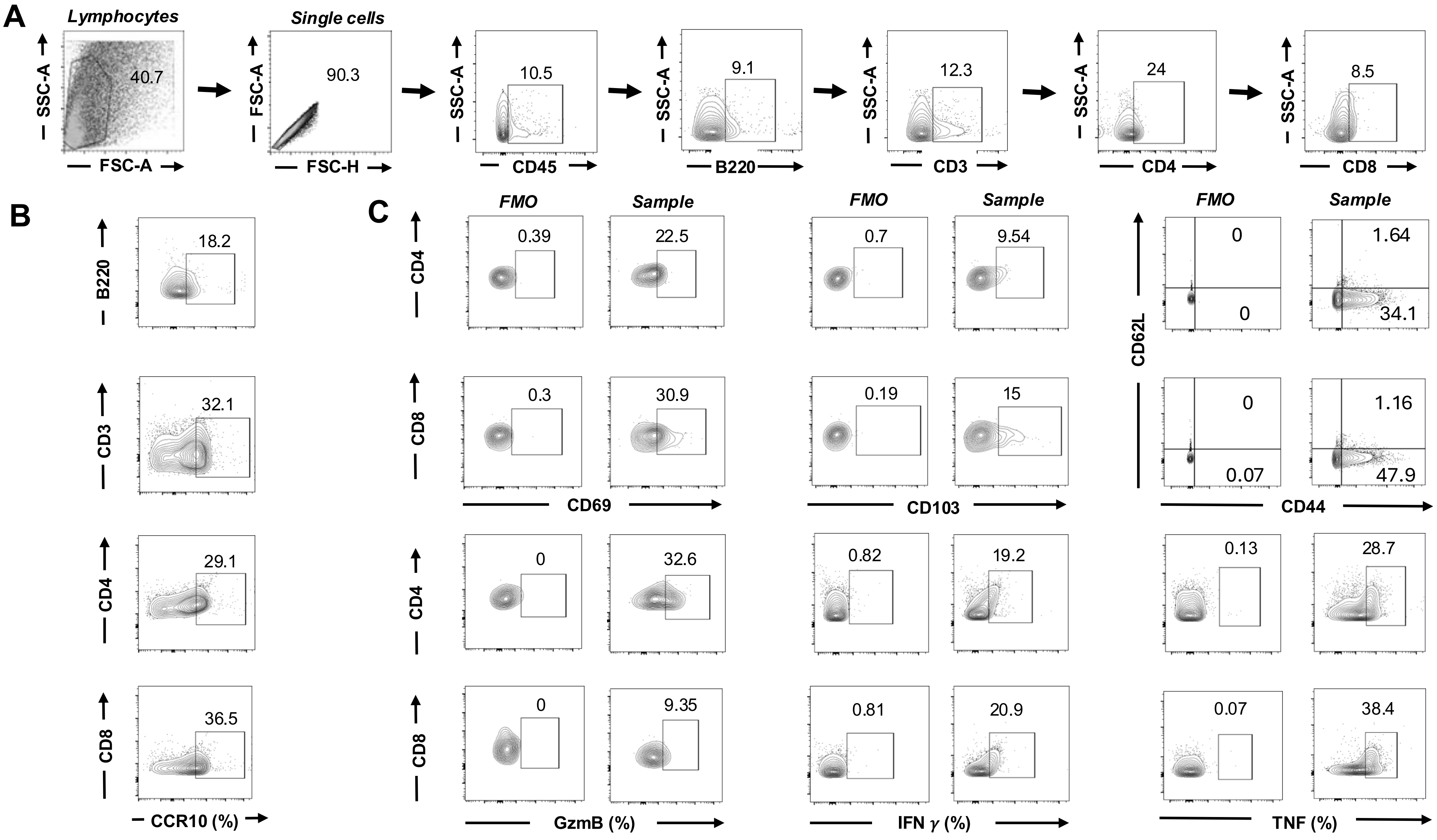
A
Lymphocytes
SSC-A
FSC-A
40.7
Single cells
FSC-A
FSC-H
90.3
SSC-A
CD45
10.5
SSC-A
B220
9.1
SSC-A
CD3
12.3
SSC-A
CD4
24
SSC-A
CD8
8.5
B
B220
18.2
CD3
32.1
CD4
29.1
CD8
36.5
CCR10 (%)
C
FMO
Sample
CD4
0.39
22.5
CD8
0.3
30.9
CD69
0.7
9.54
0.19
15
CD103
0
1.64
0
34.1
CD62L
0
1.16
0.07
47.9
CD44
CD4
0
32.6
CD8
0
9.35
GzmB (%)
0.82
19.2
0.81
20.9
IFN γ (%)
0.13
28.7
0.07
38.4
TNF (%)

**Supplementary Figure 1. Example of a flow cytometry gating strategy and phenotypic analysis of T cell and B cell subsets:** (**A**) Representative flow cytometry plots illustrating the gating hierarchy to identify lymphocyte subsets. Cells were first gated on forward scatter (FSC-A) vs. side scatter (SSC-A) to isolate the lymphocyte population, followed by singlet discrimination (FSC-H vs. FSC-A). We then gated CD45+ leukocytes to identify B cells (B220+) and T lymphocytes (CD3+ and further resolved T cells into CD4+ and CD8+ populations. (**B**) Assessment of homing receptor expression showing frequencies of $B220^+$, $CD3^+$, $CD4^+$, and $CD8^+$ expressing CCR10 (%). (**C**) Representative flow plots displaying activation and tissue-resident memory ($T_{RM}$) marker expression and naive, central memory ($T_{CM}$), and effector memory (TEM) CD69, CD103, CD62L, CD44 gated on $CD4^+$ and $CD8^+$ T cells. Intracellular cytokine and cytotoxic molecule staining of $CD4^+$ and $CD8^+$ T cells evaluating expression of Granzyme B (GzmB), Interferon-gamma (IFN-γ), and Tumor Necrosis Factor (TNF-α).
